# Determining the Minimal Background Area for Species Distribution Models: MinBar PACKAGE

**DOI:** 10.1101/571182

**Authors:** Xavier Rotllan-Puig, Anna Traveset

## Abstract

1. One of the crucial choices when modelling species distributions using pseudo-absences approaches is the delineation of the background area to fit the model. We hypothesise that there is a minimum background area around the centre of the species distribution that characterizes well enough the range of environmental conditions needed by the species to survive. Thus, fitting the model within this area should be the optimal solution in terms of both quality of the model and execution time.
2. MinBAR is an R package that calculates the optimal background area. The version 1.0.0 is implemented for MaxEnt and uses Boyce Index as a metric to assess models performance.
3. Two case studies are presented to assess the hypothesis and to illustrate the package.
4. Partial models trained with part of the species distribution often perform better than those fitted on the entire extension. MinBAR is a versatile tool that helps modellers to objectively define the optimal solution.

## Introduction

Species distribution modelling (SDM) has become an essential tool in the fields of ecology and biodiversity conservation. Its popularity, among other reasons, is due to the ease of use of software such as MaxEnt (S. J. Phillips, Anderson, & Schapire, 2006) or BIOMOD (Thuiller, Lafourcade, Engler, & Araujo, 2009), but also because of the development of the R-programming community and the free availability of biodiversity data in public repositories like GBIF (https://www.gbif.org/). However, such data is often limited to species presences only and with a lack of occurrences in poorly sampled areas. These facts limit the use of some techniques or algorithms and force to make critical assumptions and choices, which introduce different levels of uncertainty to model predictions (Jarnevich et al., 2017).

One of the crucial choices when using pseudo-absences approaches is the delineation of the background area to fit the model, also called “landscape of interest” or “study area” (Elith et al., 2011; Raes, 2012). Defining its extent, however, remains a challenge. Elith et al. (2011), for instance, argued that it has to be defined by the ecologist and limited by geographic boundaries or by how far the species can disperse. More recently, other authors have considered the interactions with other species or the sampling biases in the data set as constraints (Jarnevich et al., 2017). Yet, in many situations it is difficult to accurately define a background area, either owing to limited knowledge of the species biology or to the lack of available data (R. P. Anderson & Raza, 2010; N. Barve et al., 2011). In addition, studies are usually performed at a country or regional level and, then, the background area is constrained to an artificial or political boundary despite the species distribution might be wider (El-Gabbas & Dormann, 2018). Finally, another limitation may appear when the extent of the species is so large that it makes computations to fit the model and generate predictions highly resource-demanding and time-consuming. These limitations are particularly important when the study encompasses a high number of species with a large geographical range. Any of these situations usually lead to fit partial models, which might or might not imply a reduction of model performance (El-Gabbas & Dormann, 2018).

In this work, we hypothesize that there is a minimum background area around the centre of the species distribution (minimum buffer) that characterizes well enough the range of environmental conditions needed by the species to survive. Thus, fitting the SDM within this area should be the optimal solution in terms of both quality of the model and execution time.

### MinBAR overview

MinBAR is an R package that aims at (1) defining what is the minimum or optimal background extent necessary to fit good partial SDMs and/or (2) determining whether the background area used to fit a partial SDM is reliable enough to extract ecologically relevant conclusions from it.

### Problem

On the one hand, fitting partial SDMs might lead to underestimated predictions of species’ distribution or to biased descriptions of their niches (Sanchez-Fernandez, Lobo, & Lucia Hernandez-Manrique, 2011). On the other hand, making model calibrations and predictions of species with a large geographic range can demand a huge amount of computer resources in terms of time and memory.

To solve these problems, the idea behind the MinBAR package is to sequentially fit several concentric SDMs, with different diameter each (i.e. buffers), from the centre of the species distribution to the periphery, until a satisfactory model is reached.

### Evaluation metrics

A certain controversy exists about the best way to evaluate the performance of SDMs. One of the most widely used metrics is the AUC or area under the receiver operating characteristic (ROC) curve, although it has received several critiques because of its misuse (J. M. Lobo, Jimenez-Valverde, & Real, 2008). In particular, for the purpose of MinBAR, AUC is not the best choice because its scores are highly influenced by the defined background area, then it is only useful for assessing the performance of different models with exactly the same extent. For that reason, this package uses the Boyce Index (Hirzel, Le Lay, Helfer, Randin, & Guisan, 2006), implemented in the R package *ecospat* (Di Cola et al., 2017). However, AUC is also calculated and gathered in the outputs, although it is not used to derive conclusions.

Boyce Index (BI) is a presence-only and threshold-independent evaluator for SDMs. Among others, it is adequate in situations where the model uses background data instead of true absences, as is the case of MaxEnt (Di Cola et al., 2017). It varies between −1 and 1, where positive values indicate consistent model predictions; values close to zero indicate predictions not better than those from a random model; and negative values imply bad predictions. See Hirzel et al. (2006) and Di Cola et al. (2017) for further details on how BI is calculated as well as its strengths and weaknesses.

In order to evaluate the predictive performance of the models, this package includes two different metrics. On the one hand, Boyce Index Partial (BI_part) evaluates the accuracy of predictions within the buffer, or what is the same, in the training area. On the other hand, Boyce Index Total (BI_tot) assesses predictions beyond the training area, across the whole distribution of the species (i.e. transferability of the model).

### *minba*: The main function

The main function of MinBAR is *minba*. In the version 1.0.0 of the package presented here, *minba* is implemented for MaxEnt models.

This function firstly loads the presences’ data set and the explanatory variables. Secondly, it calculates the centre of the species distribution, the most distant occurrence and the buffers. The buffers are not defined by equal distance, but by % of presences equally distributed. This is particularly useful for very discontinuous distributions (e.g. introduced or invasive species), while not affecting more aggregated populations.

Thirdly, *minba* makes *n* models for each buffer in a loop and calculates averages. In this step, it crops the variables to the extent of the buffer +5%, and calculates the number of necessary pseudo-absences to cover the 50% of the pixels within the buffer (Guevara, Gerstner, Kass, & Anderson, 2018). It uses 70% of the presences to calibrate the model and 30% for evaluation, all from within the buffer (Boyce Index Partial). It also makes predictions and evaluations for the whole extent of the species +5% (Boyce Index Total). For this assessment, it uses 30% of all presences excluding those used to calibrate the model.

At this point, the user can choose either (1) to run the models for all the buffers to see if the selected background area is accurate and how the quality of the models evolves, or (2) to stop the process when it reaches certain conditions, which can be defined by the user as well. The latter option is adequate for very large species distributions. In this case, the user also has several options, mainly depending on the aim of the study. On the one hand, if the interest is related to the characteristics of the population (e.g. description of the ecological niche, etc.), the focus should be more in the Boyce Index Partial. On the other hand, if the aim is to project the model in time or space, the focus should fall on the Boyce Index Total. In turn, both approaches have two possibilities: either (a) fixing a minimum Boyce Index to stop the process when it is reached, or (b) to automatically stop it when the standard deviation (SD) of the last four calculated buffer’s Boyce Index is small. Thus, the user has four arguments (i.e. BI_part, BI_tot, SD_BI_part and SD_BI_tot) to pass to *minba* in order to define how to proceed. BI_part and BI_tot accept two possibilities: either *NULL* (default), which deactivates the condition, or a number below 1 (it makes no sense a higher BI), which establishes the minimum limit. Similarly, SD_BI_part and SD_BI_tot accept *NULL* (default) to deactivate the condition, or a number to establish the minimum SD. After checking the results of the case studies presented in this document (see below), a recommended minimum SD could be 0.006. Therefore, there are several combinations to choose from. For instance, if all four arguments are *NULL* (default), all buffers are modelled; alternatively, if both BI_par and BI_tot are defined as a number, and so are SD_BI_part and SD_BI_tot, the process stops when the first of them is reached. Any combination of them is allowed.

### Outputs

At the end of the modelling process, *minba* outputs different information in the form of tables and charts to let the user know the optimal buffer.

It writes out three tables in *csv* files: *selfinfo_mod_, info_mod_* and *info_mod_means_* (all followed by the name of the species). The first two tables are merely informative about how the modelling process has been developed and the results of each model. Whereas *info_mod_means_* shows the means of the *n* models run for each buffer. See Table S1 in Supplementary Material as an example of *info_mod_means_*. It contains the Boyce Index Partial, the Boyce Index Total and the execution time. Additionally, it also has columns with rankings of the buffer derived from these three metrics, plus two more ranking columns: *rankFinalNoTime* and *rankFinalWithTime*, which rank for the best buffer with and without taking into account the execution time, respectively.

Finally, *minva* draws scatterplots, smoothed by fitting a Loess regression curve, of the two BI to show the evolution of them with the increase of the buffer diameter in kilometres. It also plots the execution time by fitting a linear regression model.

### Implementation (Case Studies)

To test the hypothesis on the existence of an optimal background area, we used two different case studies. For each one we selected several common plant species of different typology (i.e. herbaceous, shrubs, broad-leaved trees, conifers). The function *minba*, by default, defines 10 buffers, with 3 model replicates per buffer, and lets the process produce models for all of them. By doing so, one can appreciate the evolution of the metrics along the different buffers. MaxEnt was run with the default parameters, except for the number of background points. The intention of that was to limit interferences in the results as much as possible for all the species. We used 19 climatic variables available from WorldClim at different resolutions for each case of study. Equally, we downloaded the occurrences of the species from public repositories by means of the PreSPickR package (Rotllan-Puig, 2018).

All the R scripts used in these case studies, including the code for the generation of the manuscript, can be found in https://github.com/xavi-rp/MinBA. In turn, the source code of the MinBAR package can be downloaded from https://github.com/xavi-rp/MinBAR.

### Case 1: Entire distribution

We modelled 25 species native from Eurasia and North of Africa (see the list in Supplementary Material Table S2.1). The presences were downloaded from GBIF. We discarded those occurrences out of the native areas as they were introduced, and this was out of the scope of this case study.

The output graphs produced by *minba* for instance for *Fraxinus excelsior* and *Linaria alpina* can be seen in Figure 1 and Figure 2, respectively. Both BI_tot and BI_part for the two species did not notably improve when increasing the buffers after the second one. A similar pattern was seen for almost all the species studied (see all plots in Supplementary Material S3). Actually, the results (Table 1, Figure 3) showed that the best models for most of the species were those fitted with only part of their distribution, both taking into account the execution time (96%) and not doing so (72%).

**Table 1:** Best and second best buffer with and without taking into account execution time, for each species in Case Study 1

| Species | Best Buffer<br>No-Time | 2nd Buffer<br>No-Time | Best Buffer<br>With-Time | 2nd Buffer<br>With-Time |
| --- | --- | --- | --- | --- |
| pin_syl | 10 | 9 | 1 | 10 |
| que_ile | 10 | 7 | 7 | 10 |
| fag_syl | 9 | 10 | 9 | 10 |
| fra_exc | 4 | 3 | 3 | 4 |
| que_pet | 9 | 8 | 9 | 8 |
| que_rob | 10 | 9 | 2 | 5 |
| que_pyr | 9 | 10 | 9 | 5 |
| que_sub | 9 | 7 | 8 | 9 |
| abi_alb | 9 | 8 | 3 | 4 |
| ace_pla | 10 | 8 | 1 | 2 |
| aln_glu | 8 | 10 | 8 | 7 |
| jun_oxy | 8 | 9 | 8 | 9 |
| arb_une | 8 | 10 | 8 | 10 |
| cra_mon | 10 | 7 | 4 | 10 |
| pru_spi | 6 | 8 | 4 | 6 |
| bux_sem | 8 | 9 | 8 | 4 |
| cot_tom | 10 | 7 | 5 | 10 |
| vio_mir | 7 | 9 | 7 | 9 |
| dip_eru | 10 | 9 | 10 | 5 |
| cen_alb | 9 | 4 | 4 | 5 |
| ger_luc | 9 | 10 | 2 | 9 |
| lin_alp | 8 | 6 | 8 | 5 |
| pis_ter | 9 | 10 | 5 | 9 |
| leo_com | 8 | 10 | 6 | 8 |
| lot_edu | 9 | 10 | 4 | 5 |

**Figure 1:**
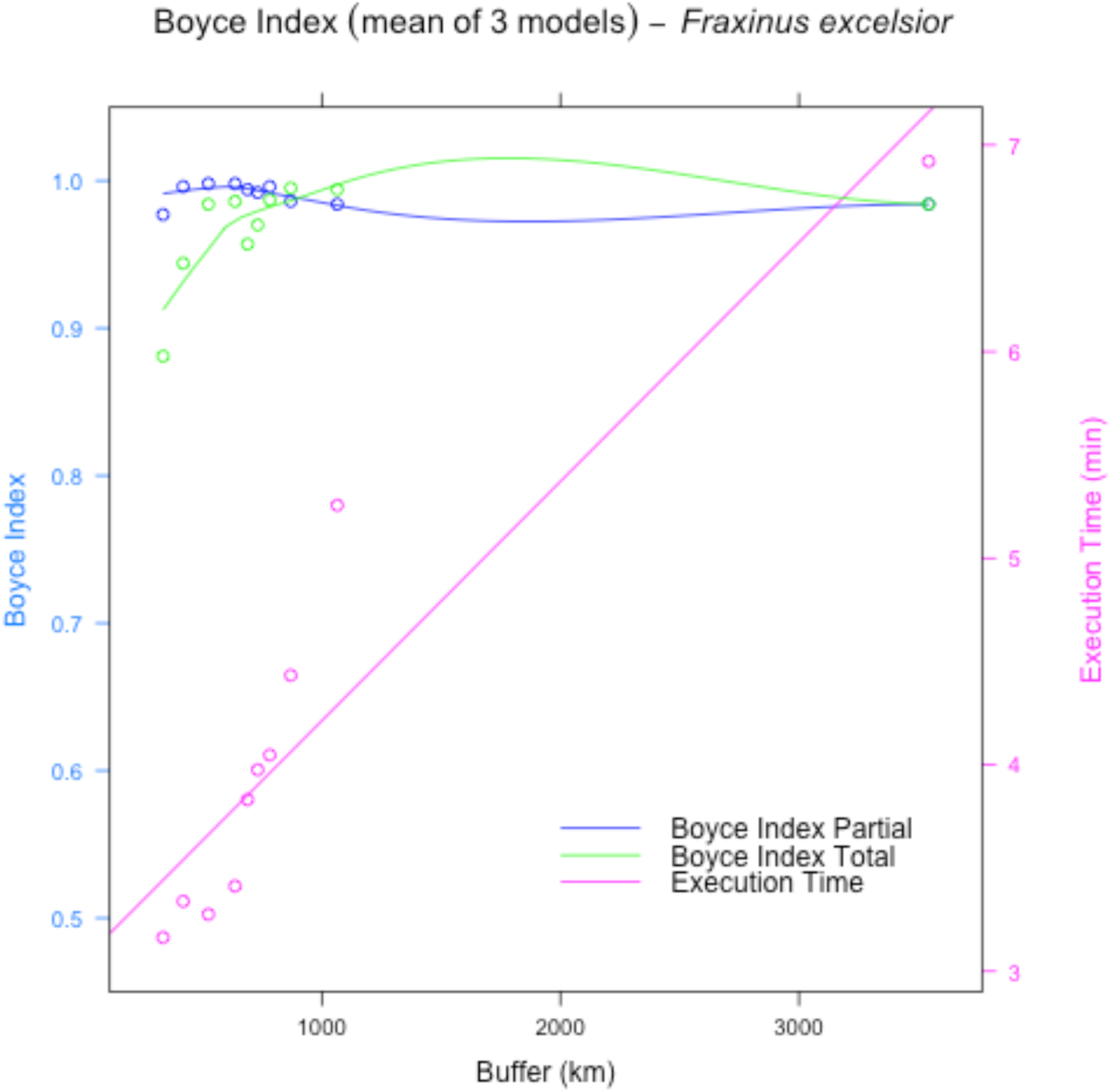
Evolution of Boyce Index Total (green) and Parcial (blue) and the execution time in minutes (pink) for Fraxinus excelsior.

**Figure 2:**
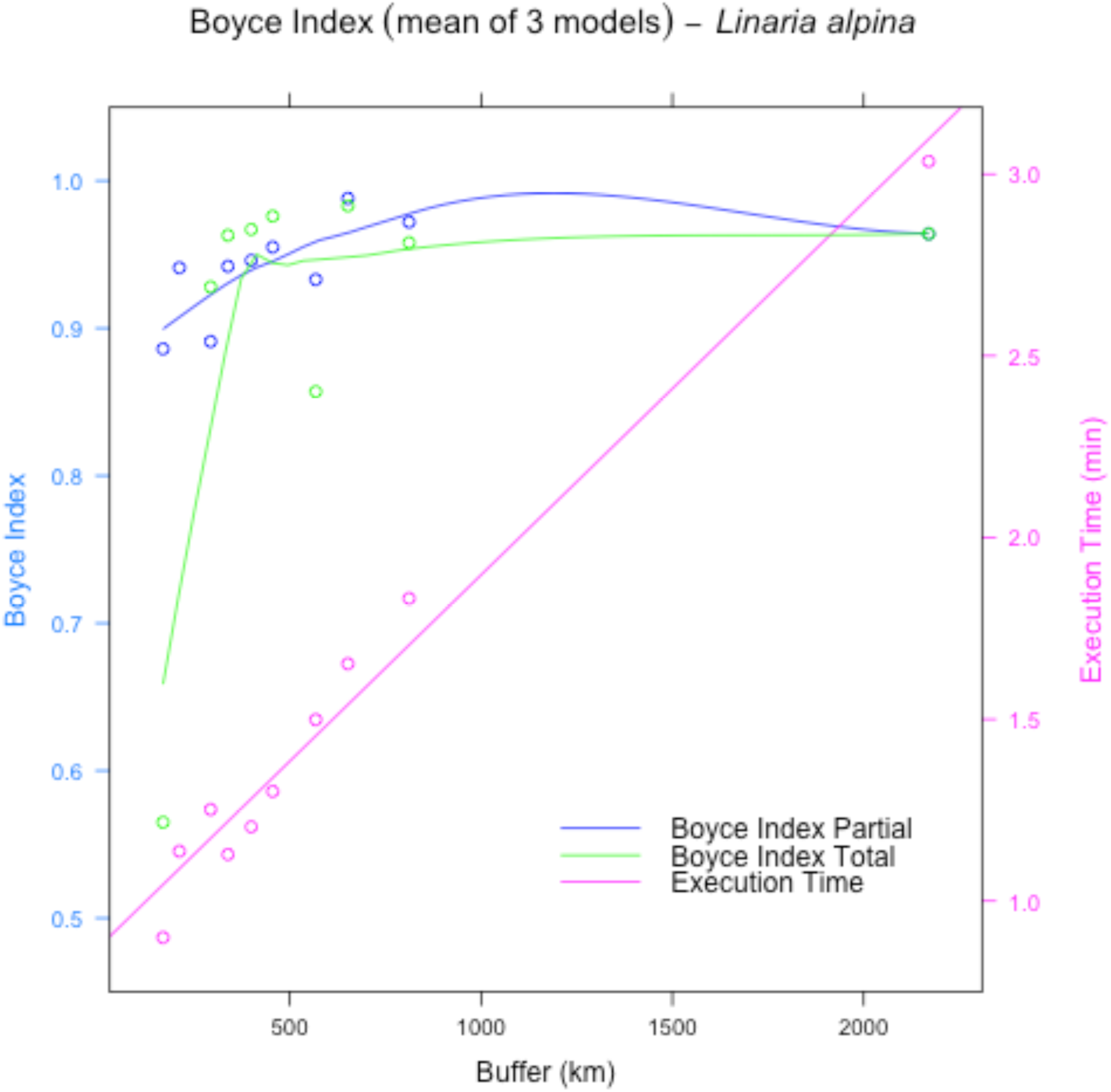
Evolution of Boyce Index Total (green) and Parcial (blue) and the execution time in minutes (pink) for Linaria alpina.

**Figure 3:**
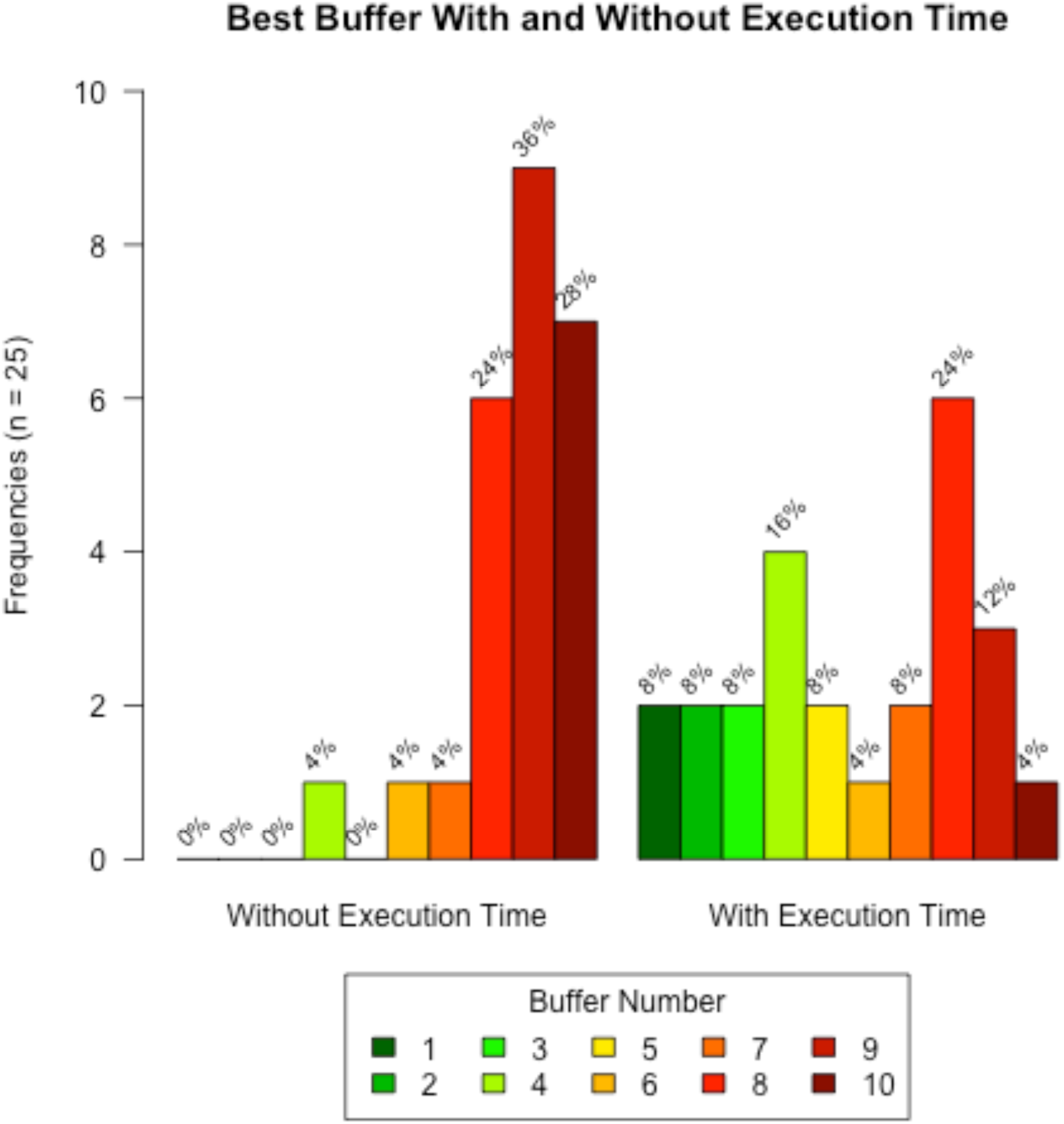
Frequencies of best buffer with and without taking into account execution time in Case Study 1.

### Case 2: Partial distribution on islands

We modelled the distribution of 10 species on the Balearic Islands (Western Mediterranean), although their native distribution also includes other continental areas (see the list in Supplementary Material Table S2.2). The occurrences were downloaded from Bioatles (http://bioatles.caib.es).

As two examples, the output graphs produced by *minba* for *Arbutus unedo* and *Asphodelus aestivus* can be seen in Figure 4 and Figure 5, respectively. Both BI_tot and BI_part for the two species did not improve very much when increasing the buffers after the first half, and a similar pattern was seen for almost all the studied species (see all plots in Supplementary Material S4). The results (Table 2, Figure 6) also showed that the best models for most of the species were those fitted with only part of their distribution, specially taking into account the execution time (90%) but also not doing so (70%).

**Table 2:** Best and second best buffer with and without taking into account execution time, for each species in Case Study 2

| Species | Best Buffer | 2nd Buffer | Best Buffer | 2nd Buffer |
| --- | --- | --- | --- | --- |
|  | No-Time | No-Time | With-Time | With-Time |
| arb_une | 8 | 3 | 8 | 3 |
| asp_aes | 10 | 8 | 1 | 2 |
| cha_hum | 5 | 7 | 5 | 2 |
| eph_fra | 8 | 10 | 6 | 8 |
| hel_sto | 9 | 6 | 6 | 9 |
| jun_oxy | 4 | 3 | 3 | 4 |
| pis_len | 10 | 9 | 10 | 3 |
| que_coc | 6 | 7 | 6 | 5 |
| rha_ala | 10 | 6 | 2 | 6 |
| vib_tin | 7 | 5 | 5 | 7 |

**Figure 4:**
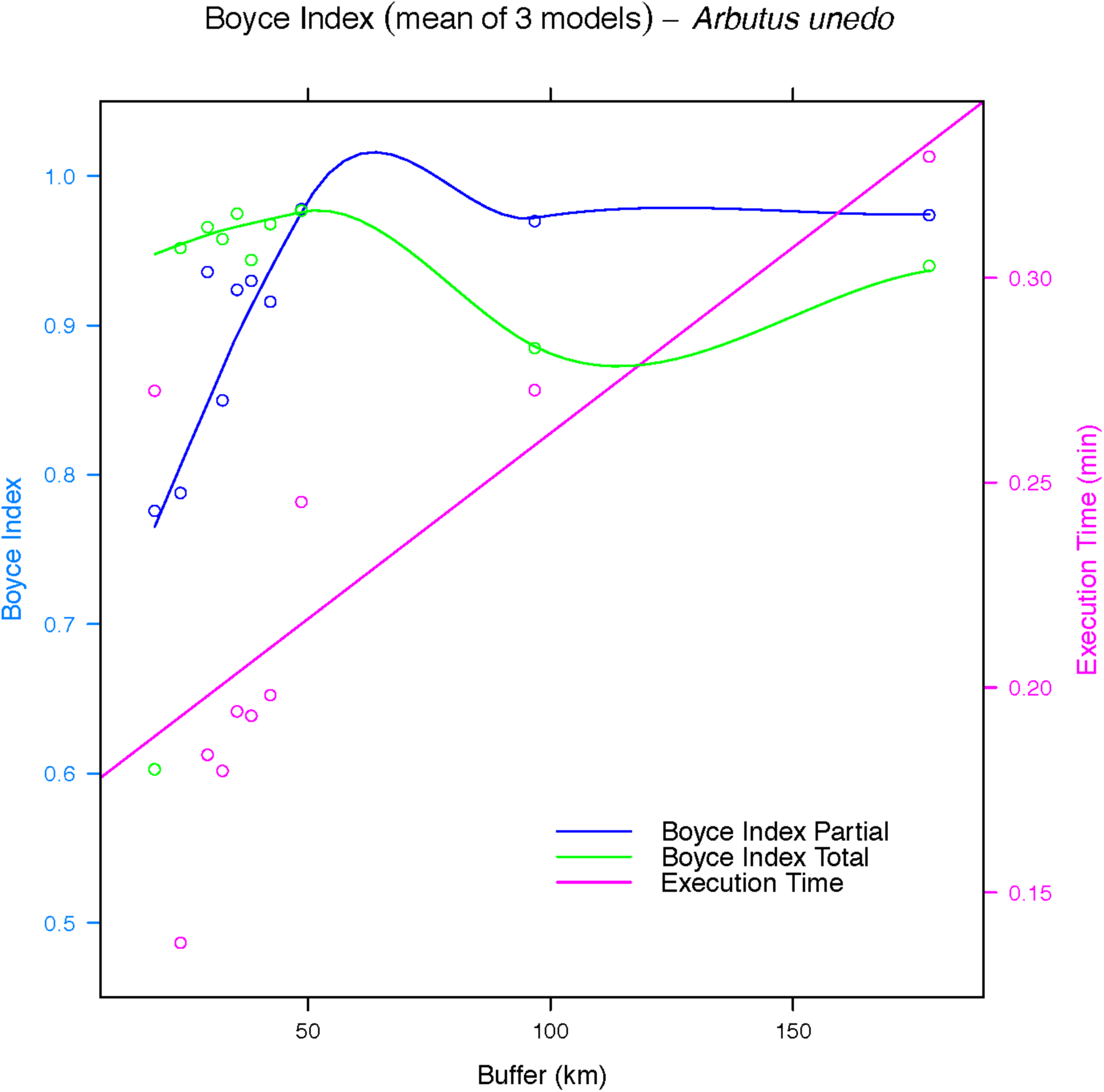
Evolution of Boyce Index Total (green) and Parcial (blue) and the execution time in minutes (pink) for Arbutus unedo.

**Figure 5:**
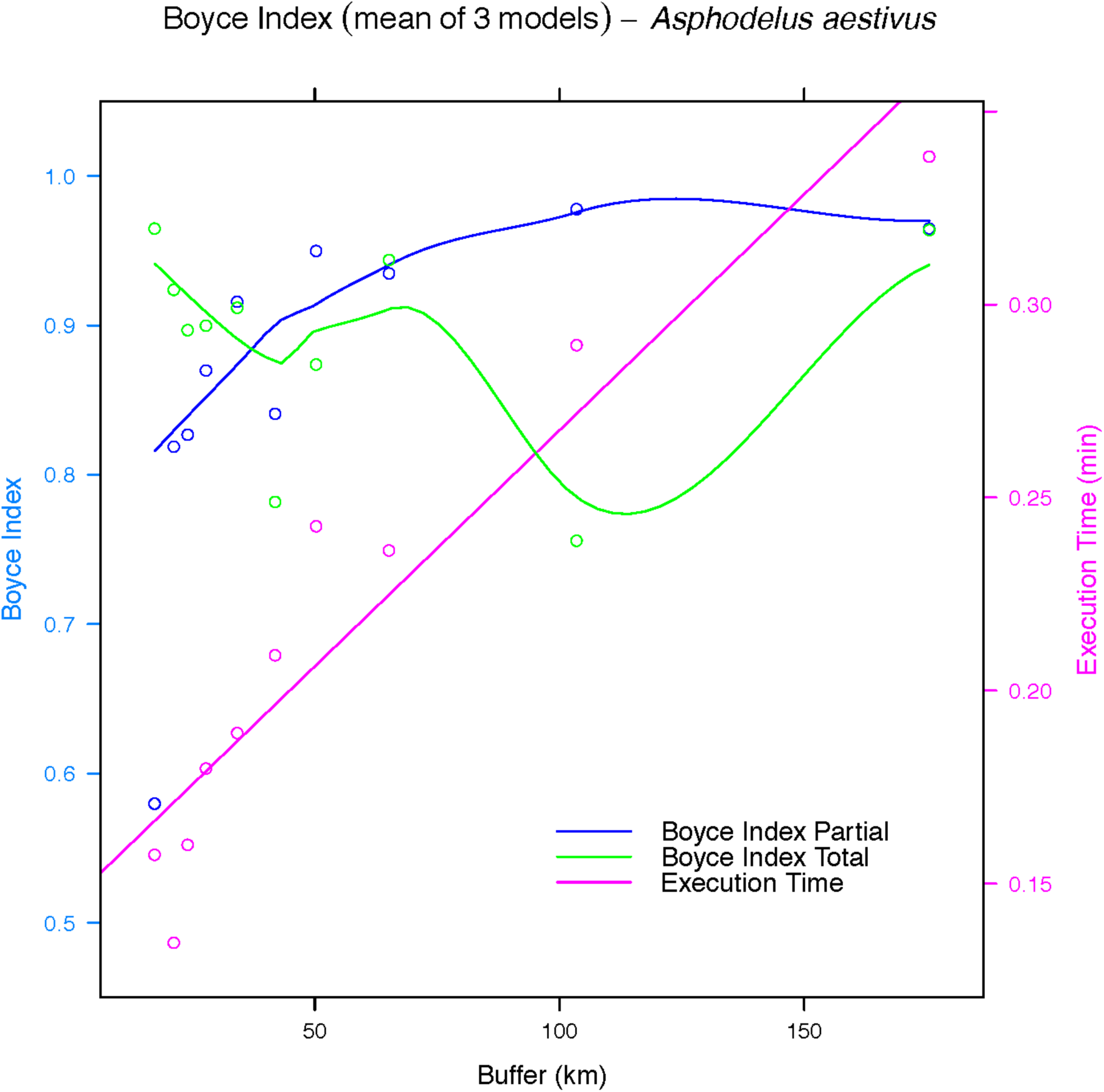
Evolution of Boyce Index Total (green) and Parcial (blue) and the execution time in minutes (pink) for Asphodelus aestivus.

**Figure 6:**
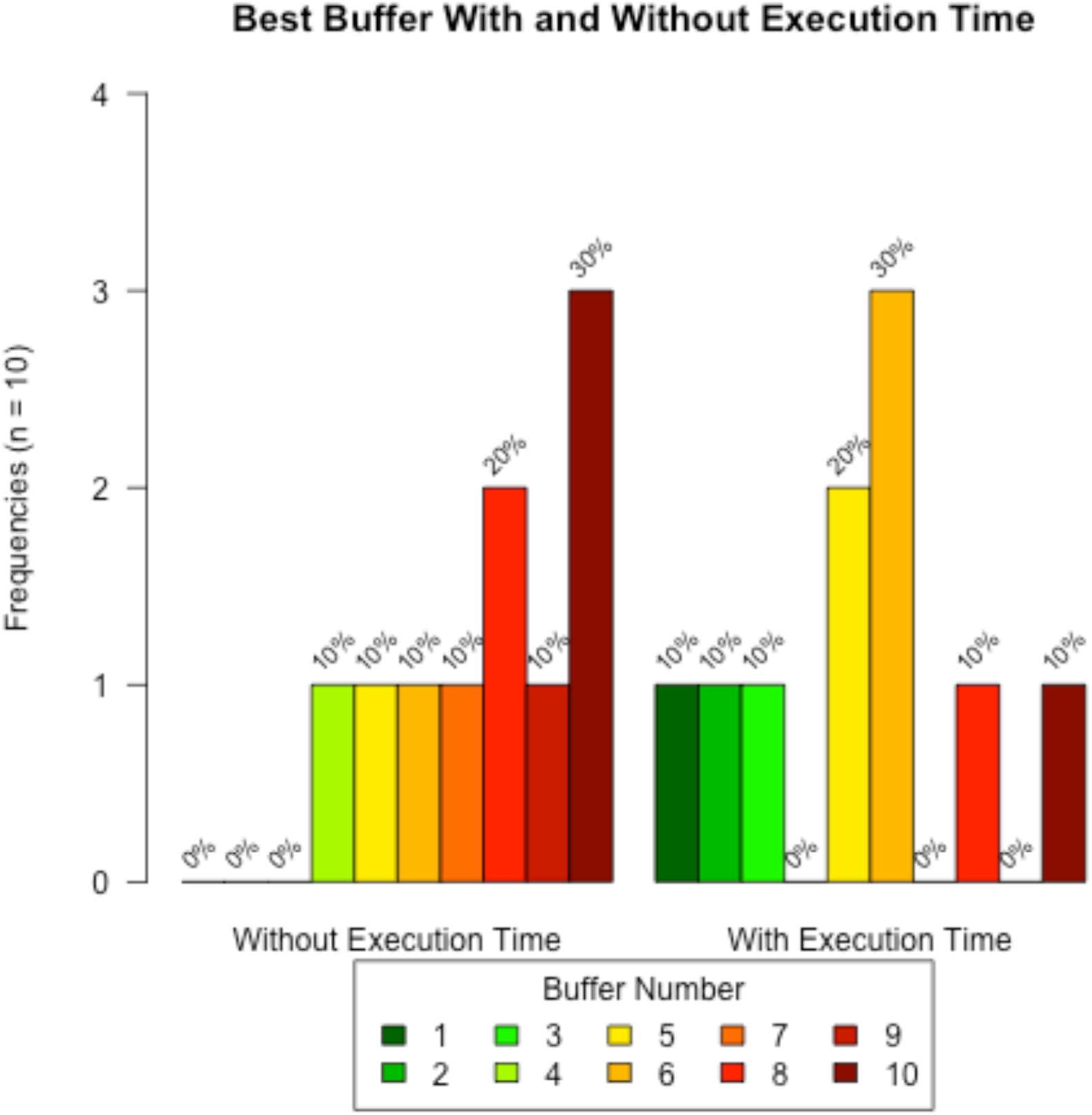
Frequencies of best buffer with and without taking into account execution time in Case Study 2.

## Conclusions

The package MinBAR has been developed, so far, to work with MaxEnt. It includes the Boyce Index as the main evaluator of the models predictive performance. In coming versions, however, it would be interesting to include other threshold-dependent evaluators based on sensitivity and specificity, as well as the option to pass arguments to the *maxent* function, or to decide the centre from where to start delimiting buffers for modelling. In addition, the inclusion of an index that would take into account at the same time the accuracy in the training area and after transferring to further areas, such as the one described by Duque-Lazo et al. (2016), might also be quite useful for the users. Furthermore, the implementation of other algorithms and modelling techniques would be highly convenient.

In short, delimiting the background area can strongly affect the results of SDMs (Acevedo, Jimenez-Valverde, Lobo, & Real, 2012). Both case studies presented here show that the model including the presences from all the species distribution does not always perform the best. Therefore, the tool developed here will help modellers to objectively define an optimal solution.

