## Supplementary Material for "Determining the Minimal Background Area for Species Distribution Models: MinBar PACKAGE"

### Introduction

This is the Supplementary Material to the article “Determining the minimal background area for MaxEnt species distribution models: MinBAR package” (https://github.com/xavi-rp/MinBAR)

### Supplementary Material S1

*Table S1: Example of an output of MinBAR. Buffer in km. (continued below)*

| Species | Buffer | BoyceIndex_part | BoyceIndex_tot | SD_part |
|---|---|---|---|---|
| Prunus spinosa | 126.8 | 0.986 | 0.843 | NA |
| Prunus spinosa | 228.1 | 0.992 | 0.911 | NA |
| Prunus spinosa | 304.2 | 0.988 | 0.913 | NA |
| Prunus spinosa | 384.3 | 0.999 | 0.946 | 0.005737 |
| Prunus spinosa | 476.5 | 0.998 | 0.912 | 0.005188 |
| Prunus spinosa | 591.5 | 0.999 | 0.988 | 0.005354 |
| Prunus spinosa | 746.2 | 0.988 | 0.976 | 0.005354 |
| Prunus spinosa | 878.1 | 0.999 | 0.995 | 0.005354 |
| Prunus spinosa | 1068 | 0.998 | 0.98 | 0.005354 |
| Prunus spinosa | 4942 | 0.999 | 0.999 | 0.005354 |

*Table continues below*

| SD_tot | ExecutionTime | rankBI_part | rankBI_tot | rankTime |
|---|---|---|---|---|
| NA | 3.039 | 10 | 10 | 1 |
| NA | 3.461 | 7 | 9 | 2 |
| NA | 3.655 | 8 | 7 | 3 |
| 0.04325 | 3.794 | 1 | 6 | 4 |
| 0.01702 | 3.814 | 5 | 8 | 5 |

| | | | | |
|---|---|---|---|---|
| 0.03584 | 4.017 | 2 | 3 | 6 |
| 0.03396 | 4.29 | 9 | 5 | 7 |
| 0.03799 | 5.061 | 3 | 2 | 9 |
| 0.008461 | 5.048 | 6 | 4 | 8 |
| 0.01121 | 8.097 | 4 | 1 | 10 |

| rankFinalNoTime | rankFinalWithTime |
|---|---|
| 10 | 9 |
| 9 | 5 |
| 8 | 6 |
| 4 | 1 |
| 6 | 7 |
| 1 | 2 |
| 7 | 10 |
| 2 | 3 |
| 5 | 8 |
| 3 | 4 |

### Supplementary Material S2

*Table S2.1: List of species used in case study 1*

| Case.Study.1 | Abbreviation1 |
|---|---|
| Pinus sylvestris L. | pin_syl |
| Quercus ilex L. | que_ile |
| Fagus sylvatica L. | fag_syl |
| Fraxinus excelsior L. | fra_exc |
| Quercus petraea (Matt.) Liebl. | que_pet |
| Quercus robur L. | que_rob |
| Quercus pyrenaica Willd. | que_pyr |
| Quercus suber L. | que_sub |
| Abies alba Mill. | abi_alb |
| Acer platanoides L. | ace_pla |
| Alnus glutinosa (L.) Gaertn. | aln_glu |
| Juniperus oxycedrus L. | jun_oxy |
| Arbutus unedo L. | arb_une |
| Crataegus monogyna Jacq. | cra_mon |
| Prunus spinosa L. | pru_spi |
| Buxus sempervirens L. | bux_sem |
| Cotoneaster tomentosus Lindl. | cot_tom |
| Viola mirabilis L. | vio_mir |
| Diplotaxis erucoides DC. | dip_eru |
| Centaurea alba L. | cen_alb |
| Geranium lucidum L. | ger_luc |
| Linaria alpina Mill. | lin_alp |

| | |
|---|---|
| Pistacia terebinthus L. | pis_ter |
| Muscari comosum (L.) Mill. | leo_com |
| Lotus edulis L. | lot_edu |

*Table S2.2: List of species used in case study 2*

| Case.Study.2 | Abbreviation2 |
|---|---|
| Arbutus unedo L. | arb_une |
| Asphodelus aestivus Rchb. | asp_aes |
| Chamaerops humilis L. | cha_hum |
| Ephedra fragilis subsp. fragilis Desf. | eph_fra |
| Helichrysum stoechas (L.) Moench | hel_sto |
| Juniperus oxycedrus subsp. oxycedrus L. | jun_oxy |
| Pistacia lentiscus L. | pis_len |
| Quercus coccifera L. | que_coc |
| Rhamnus alaternus L. | rha_ala |
| Viburnum tinus L. | vib_tin |

### Supplementary Material S3

Figures S3.1 - S3.25: Evolution of Boyce Index Total (green) and Parcial (blue), and the execution time in minutes (pink), for all the species in Case Study 1

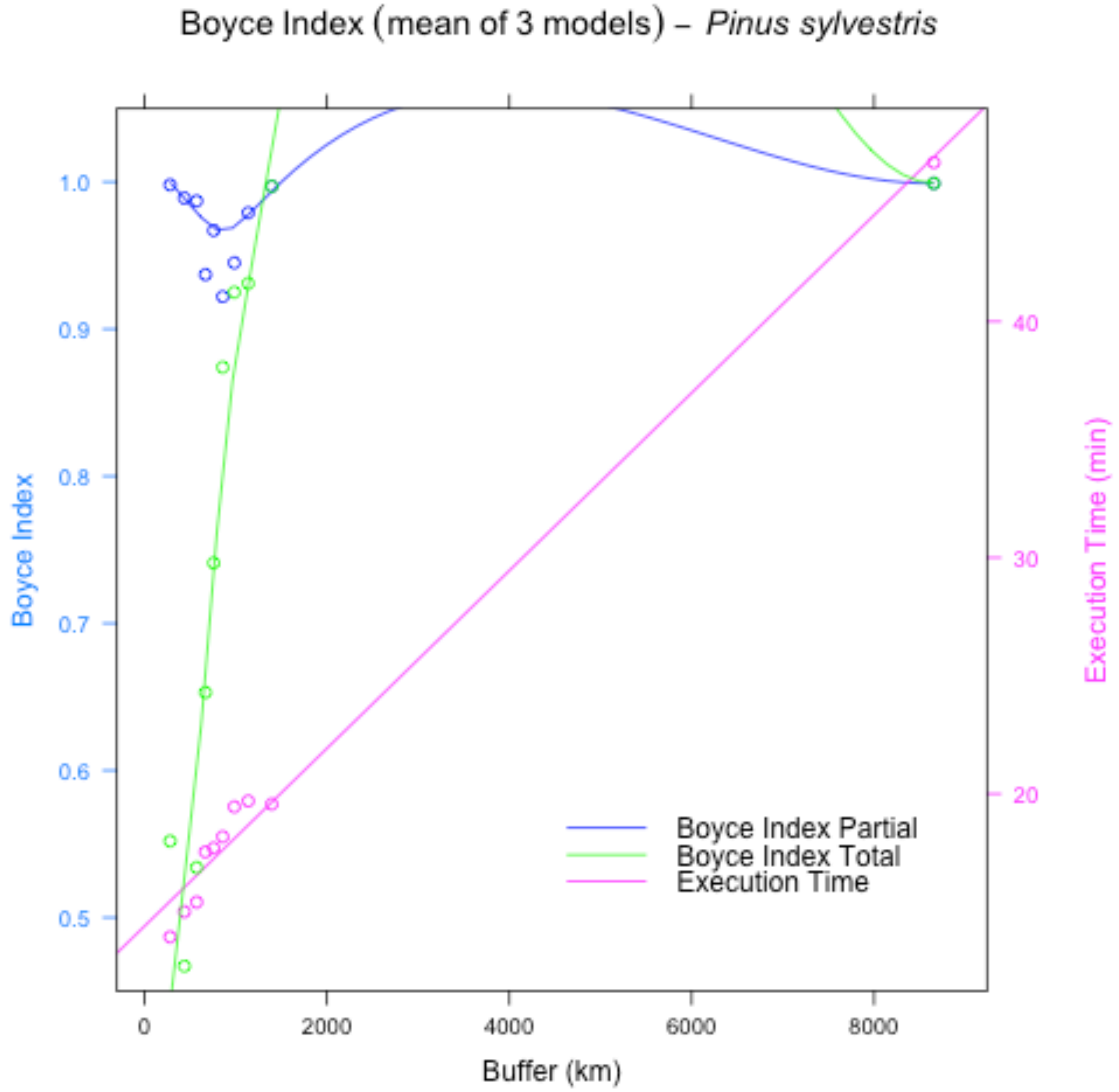

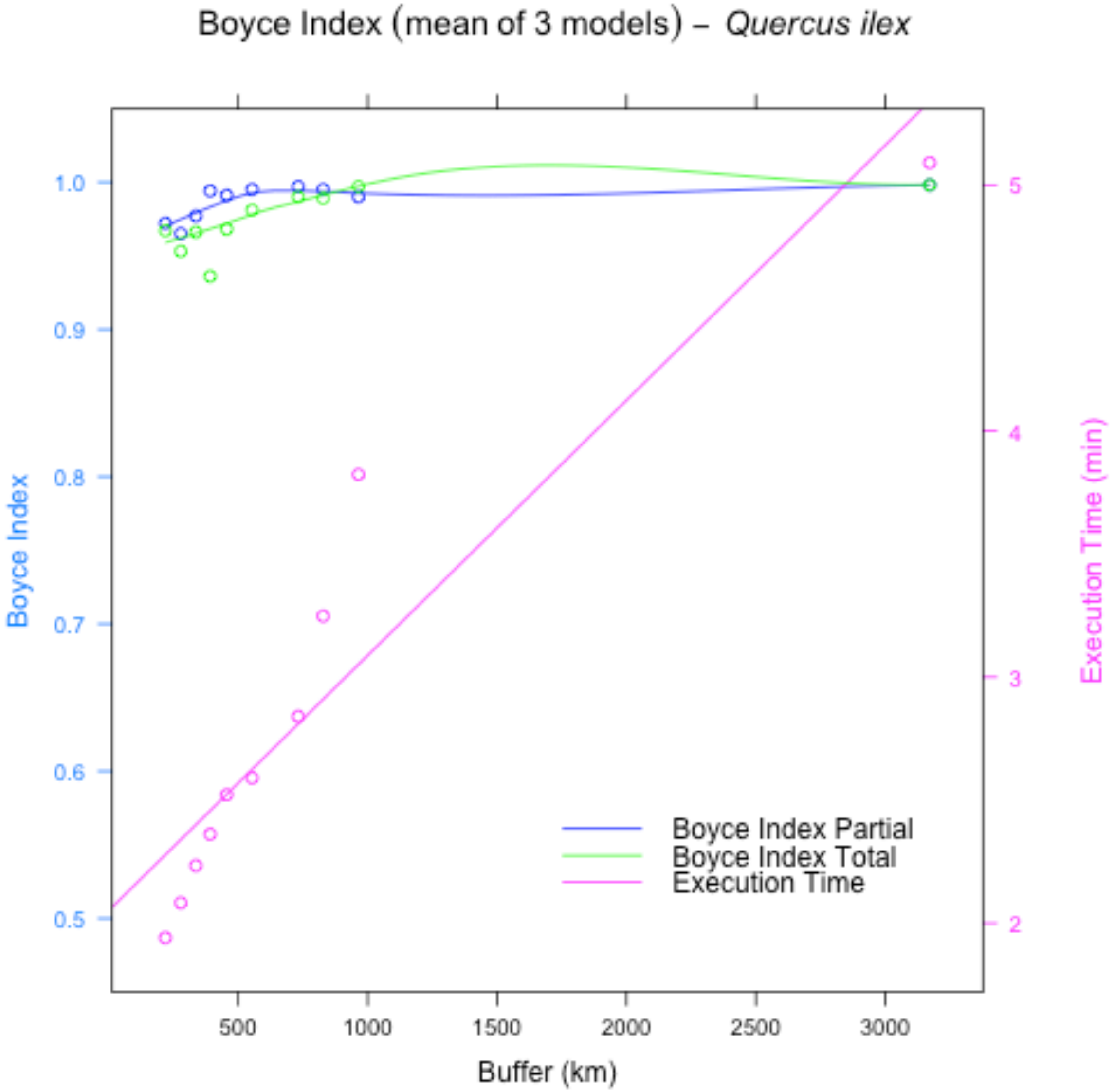
Boyce Index (mean of 3 models) – Quercus ilex
Boyce Index
1.0
0.9
0.8
0.7
0.6
0.5
Execution Time (min)
5
4
3
2
500
1000
1500
2000
2500
3000
Buffer (km)
Boyce Index Partial
Boyce Index Total
Execution Time

32

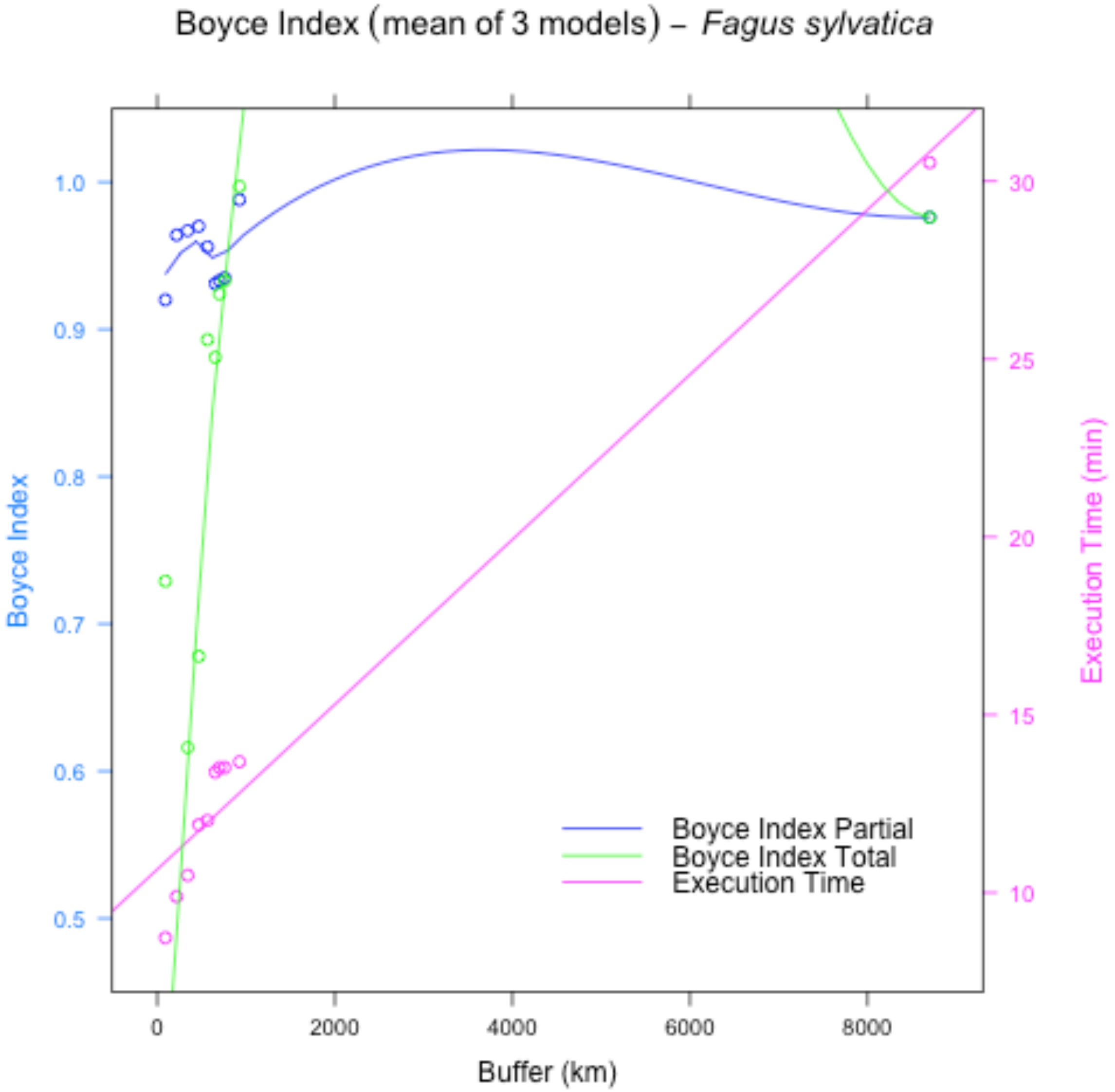

Boyce Index (mean of 3 models) – *Fagus sylvatica*
Boyce Index
1.0
0.9
0.8
0.7
0.6
0.5
Execution Time (min)
30
25
20
15
10
0
2000
4000
6000
8000
Buffer (km)
Boyce Index Partial
Boyce Index Total
Execution Time


33

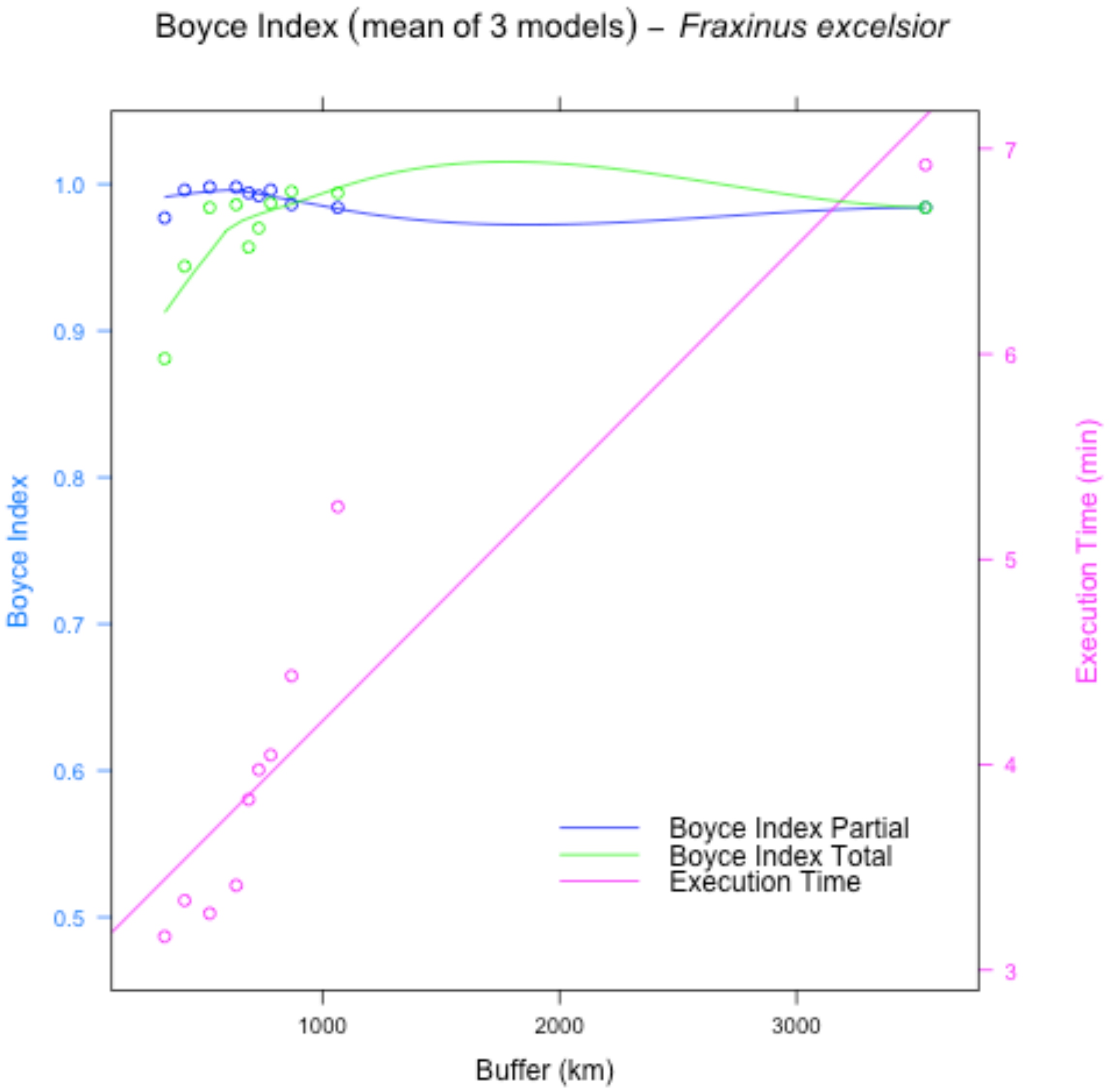
Boyce Index (mean of 3 models) – *Fraxinus excelsior*
Boyce Index
1.0
0.9
0.8
0.7
0.6
0.5
Execution Time (min)
7
6
5
4
3
1000
2000
3000
Buffer (km)
Boyce Index Partial
Boyce Index Total
Execution Time

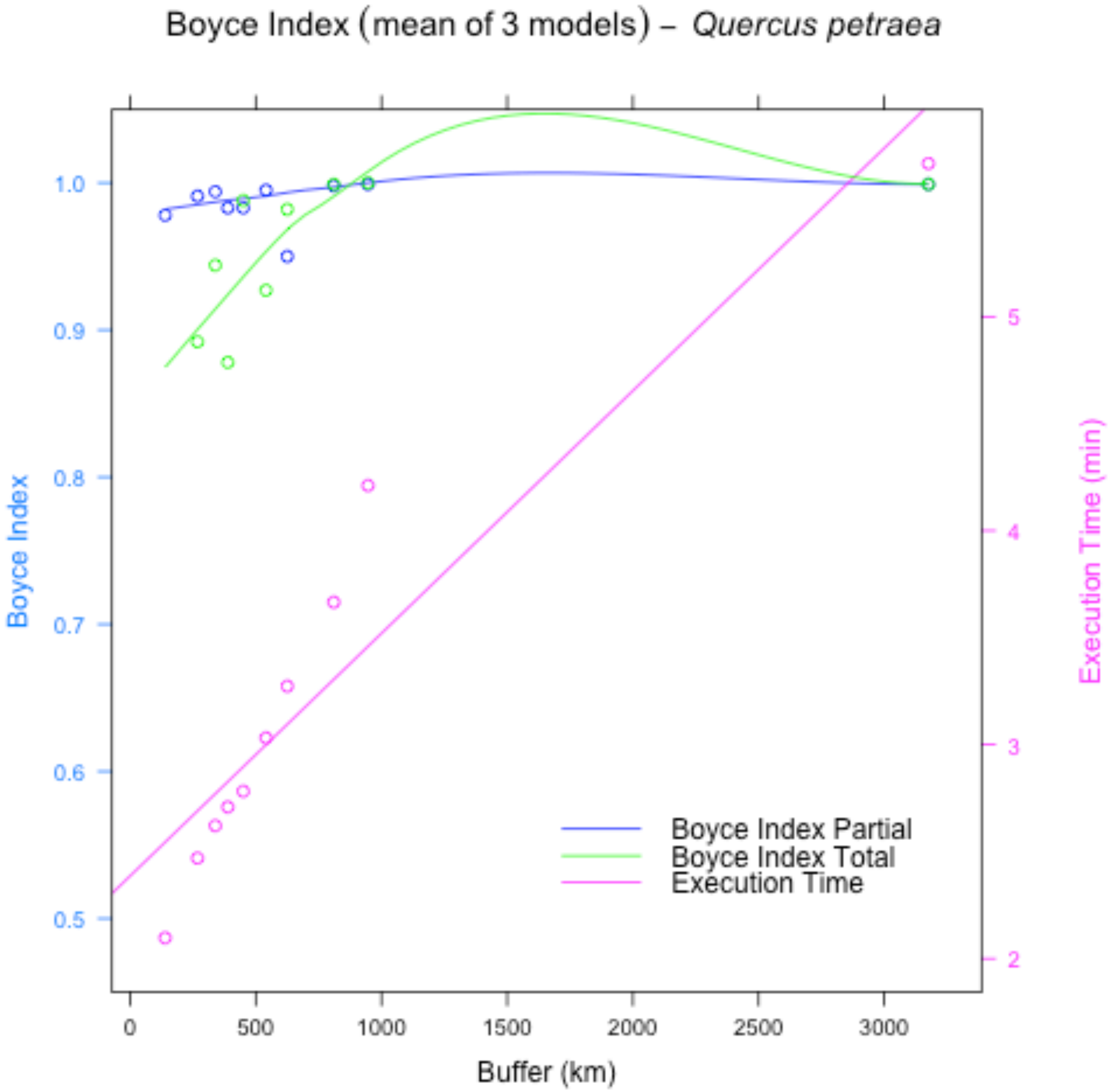
Boyce Index (mean of 3 models) – Quercus petraea
Boyce Index
Execution Time (min)
Buffer (km)
Boyce Index Partial
Boyce Index Total
Execution Time
0
500
1000
1500
2000
2500
3000
0.5
0.6
0.7
0.8
0.9
1.0
2
3
4
5

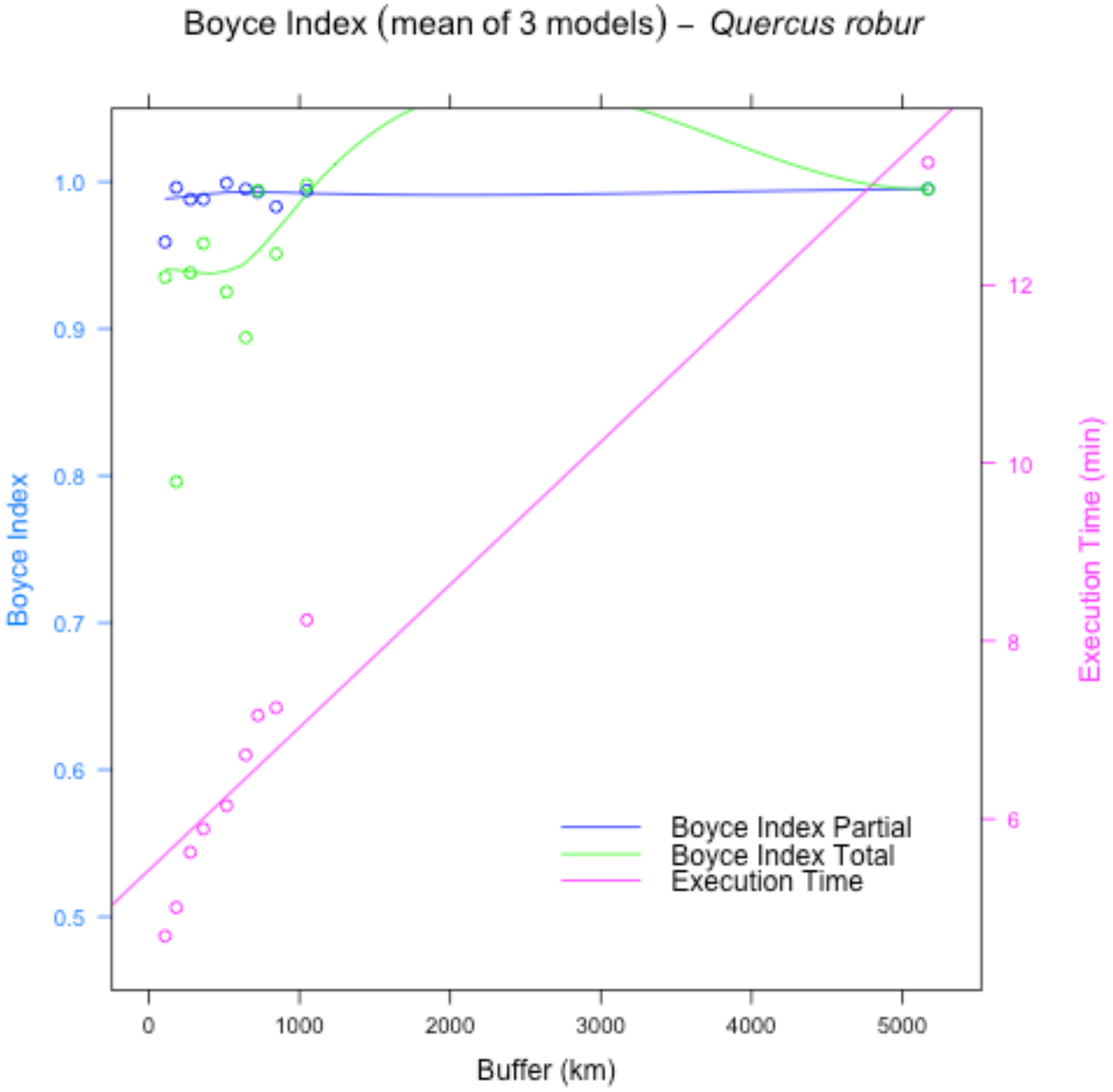
Boyce Index (mean of 3 models) – Quercus robur
Boyce Index
1.0
0.9
0.8
0.7
0.6
0.5
Execution Time (min)
12
10
8
6
0
1000
2000
3000
4000
5000
Buffer (km)
Boyce Index Partial
Boyce Index Total
Execution Time

36

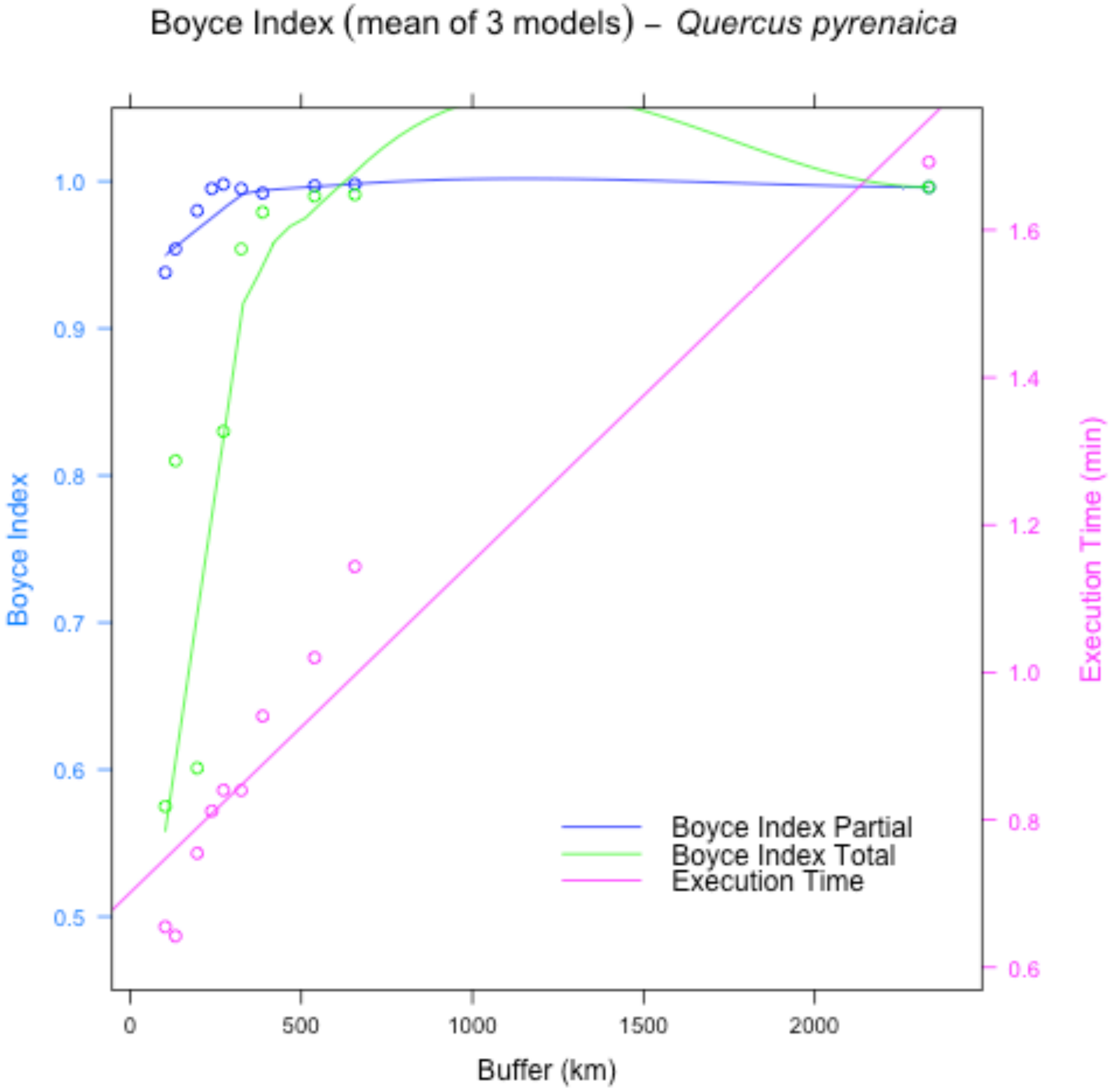
Boyce Index (mean of 3 models) – *Quercus pyrenaica*
Boyce Index
Execution Time (min)
Buffer (km)
Boyce Index Partial
Boyce Index Total
Execution Time
0.5
0.6
0.7
0.8
0.9
1.0
0.6
0.8
1.0
1.2
1.4
1.6
0
500
1000
1500
2000

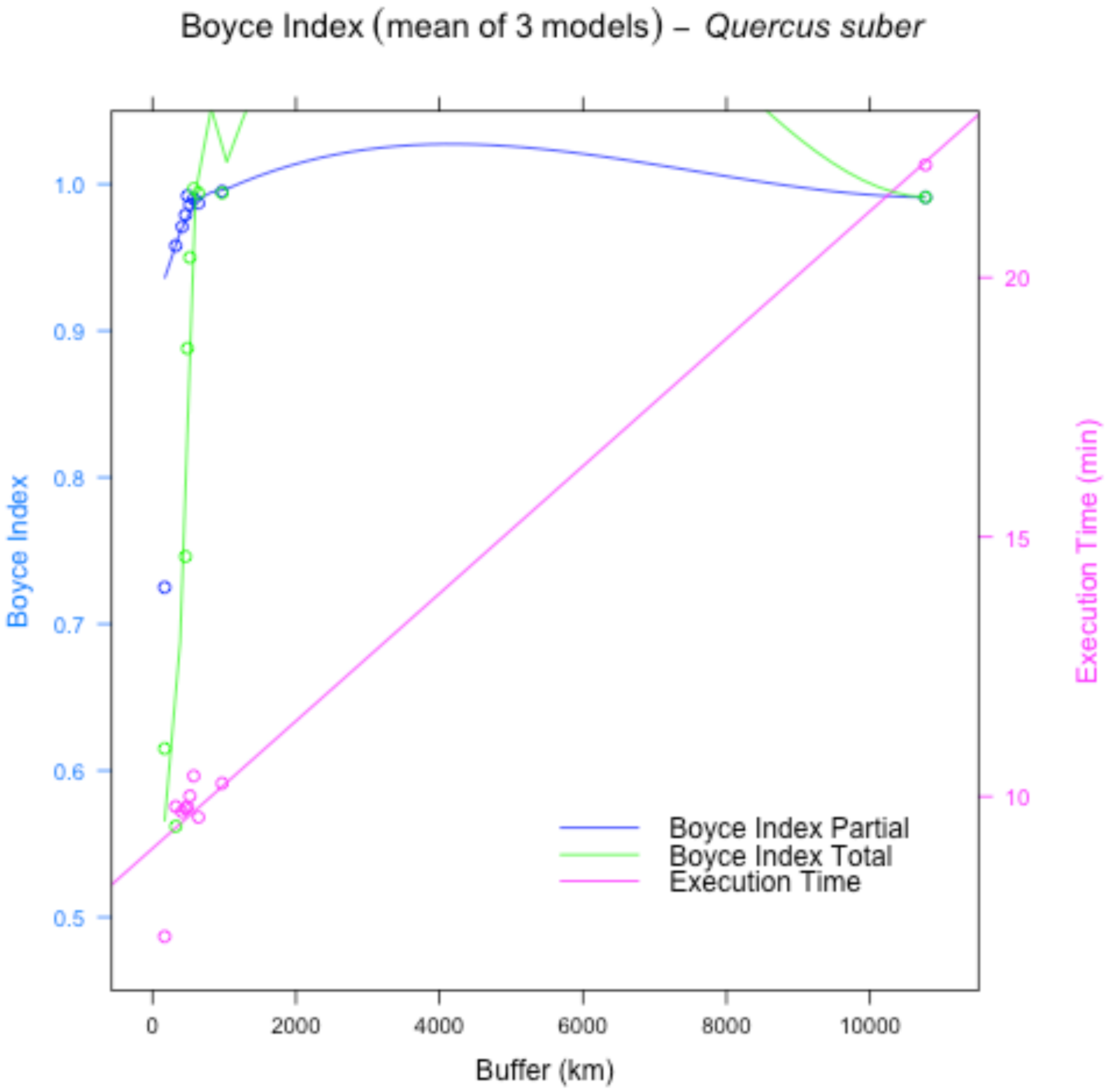
Boyce Index (mean of 3 models) – Quercus suber
Boyce Index
Execution Time (min)
Buffer (km)
0
2000
4000
6000
8000
10000
0.5
0.6
0.7
0.8
0.9
1.0
10
15
20
Boyce Index Partial
Boyce Index Total
Execution Time

38

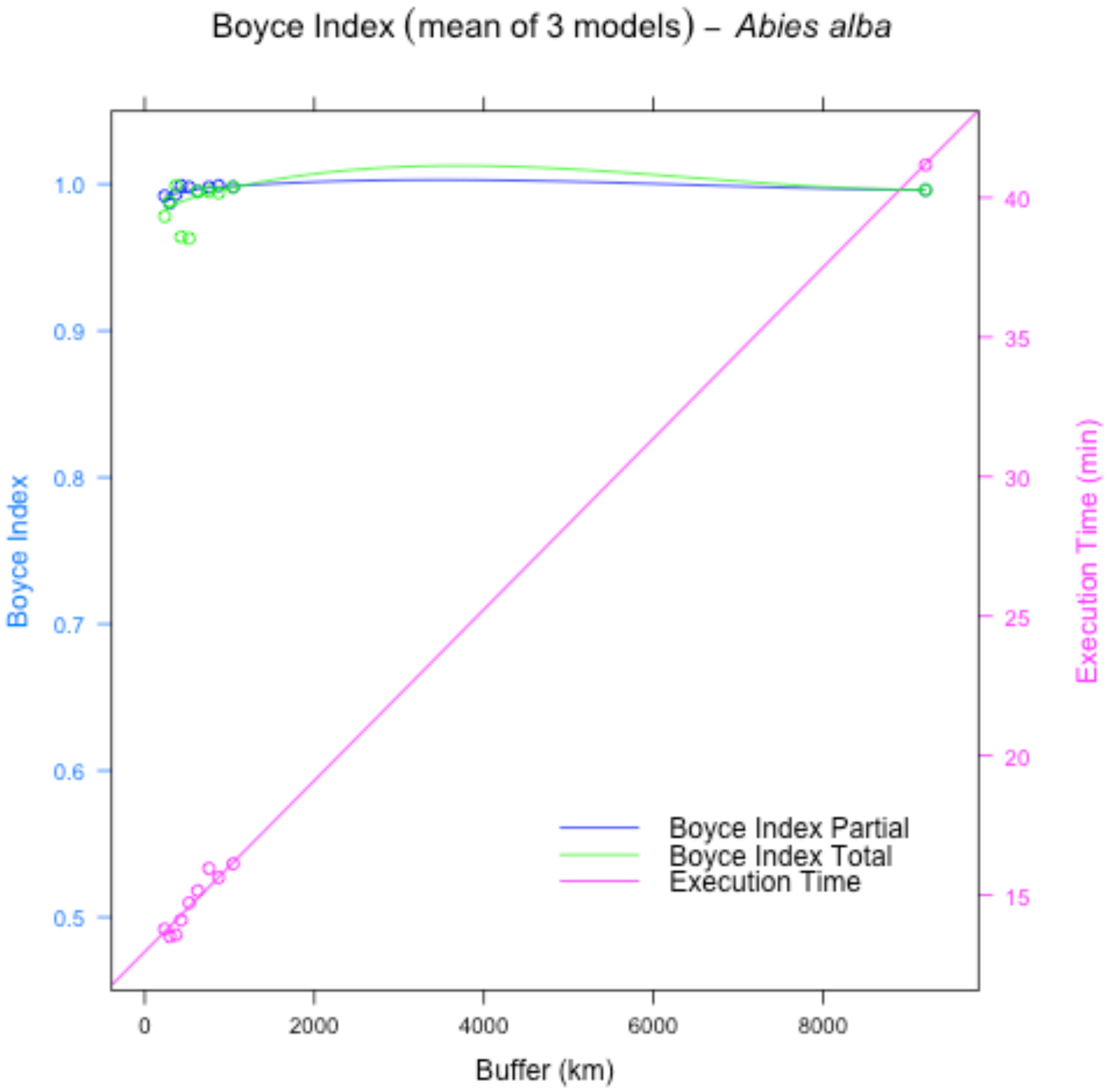

Boyce Index (mean of 3 models) – *Abies alba*
Boyce Index
Execution Time (min)
Buffer (km)
0
2000
4000
6000
8000
0.5
0.6
0.7
0.8
0.9
1.0
15
20
25
30
35
40
Boyce Index Partial
Boyce Index Total
Execution Time


39

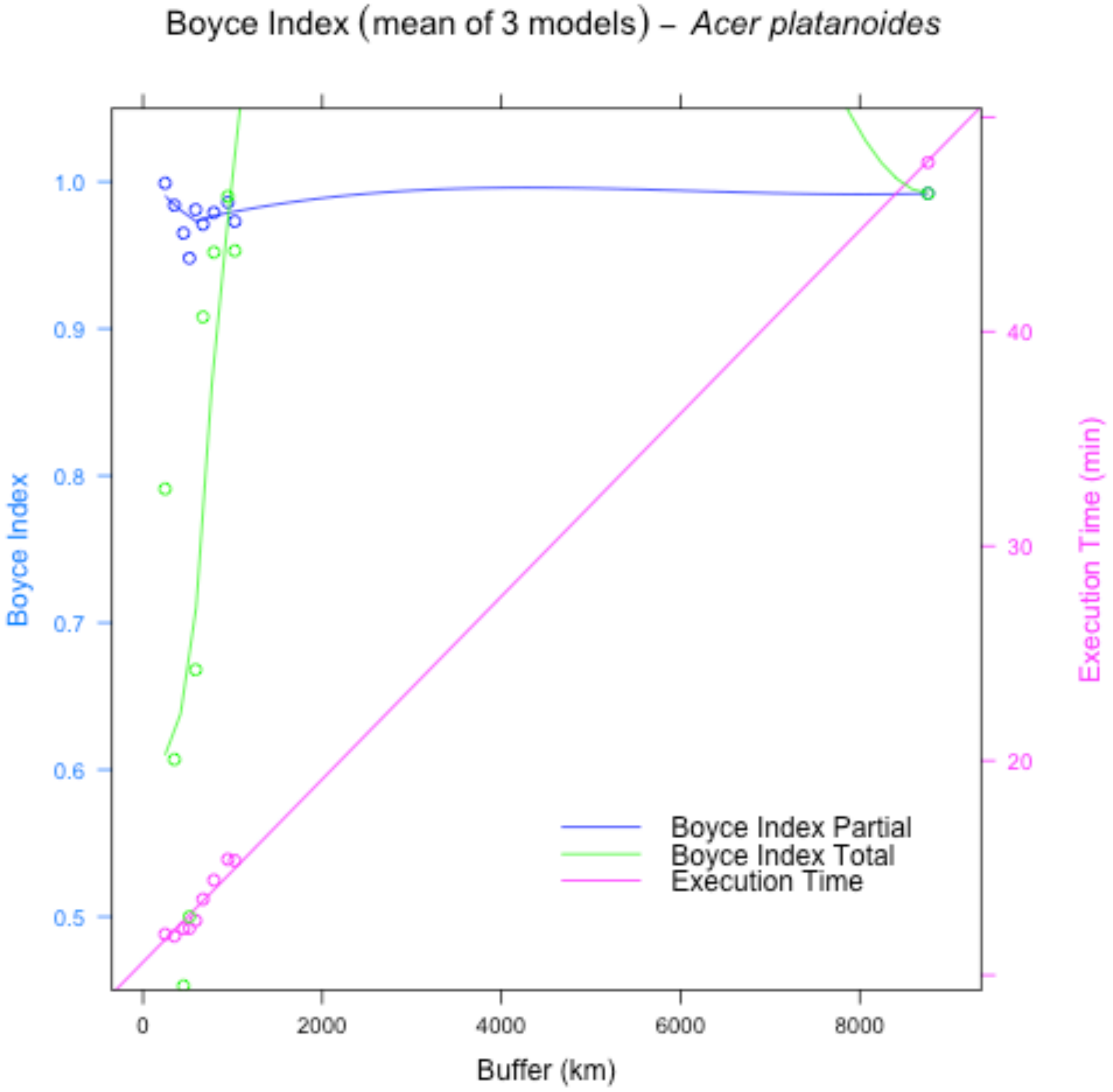
Boyce Index (mean of 3 models) – *Acer platanoides*
Boyce Index
1.0
0.9
0.8
0.7
0.6
0.5
Execution Time (min)
40
30
20
0
2000
4000
6000
8000
Buffer (km)
Boyce Index Partial
Boyce Index Total
Execution Time

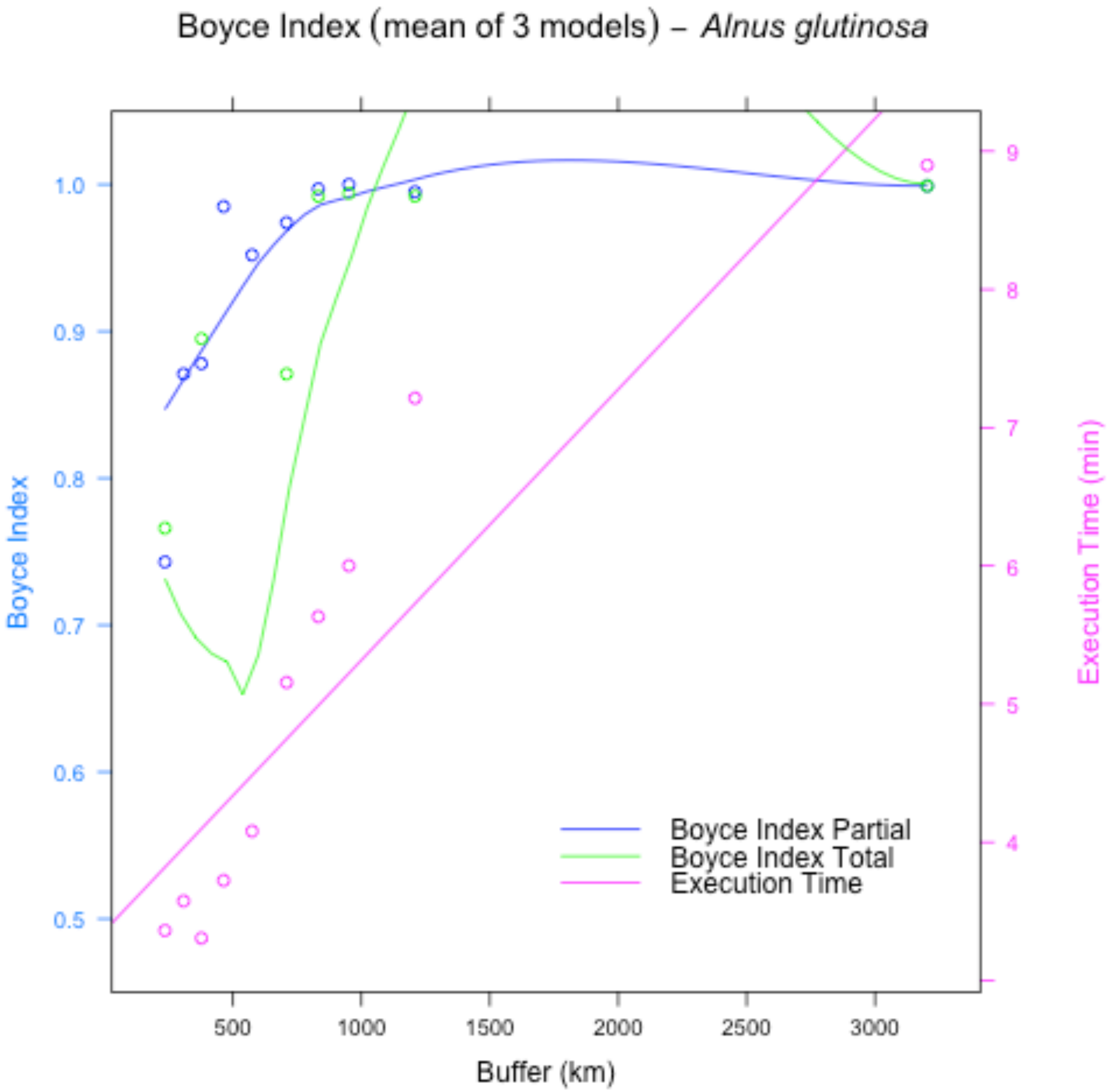
Boyce Index (mean of 3 models) – *Alnus glutinosa*
Boyce Index
Execution Time (min)
Buffer (km)
1.0
0.9
0.8
0.7
0.6
0.5
9
8
7
6
5
4
500
1000
1500
2000
2500
3000
Boyce Index Partial
Boyce Index Total
Execution Time

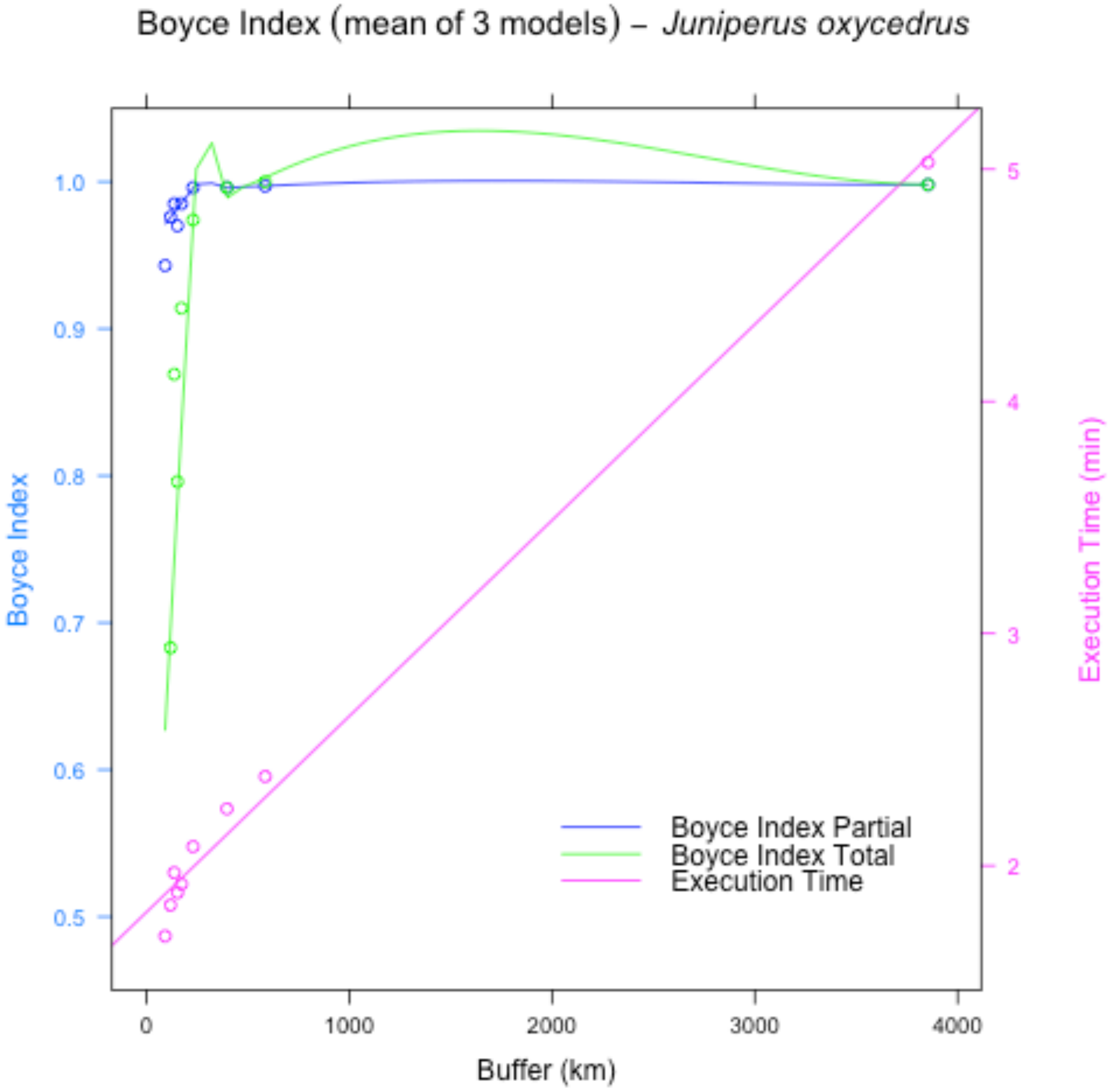
Boyce Index (mean of 3 models) – Juniperus oxycedrus
Boyce Index
Execution Time (min)
Buffer (km)
0
1000
2000
3000
4000
0.5
0.6
0.7
0.8
0.9
1.0
2
3
4
5
Boyce Index Partial
Boyce Index Total
Execution Time

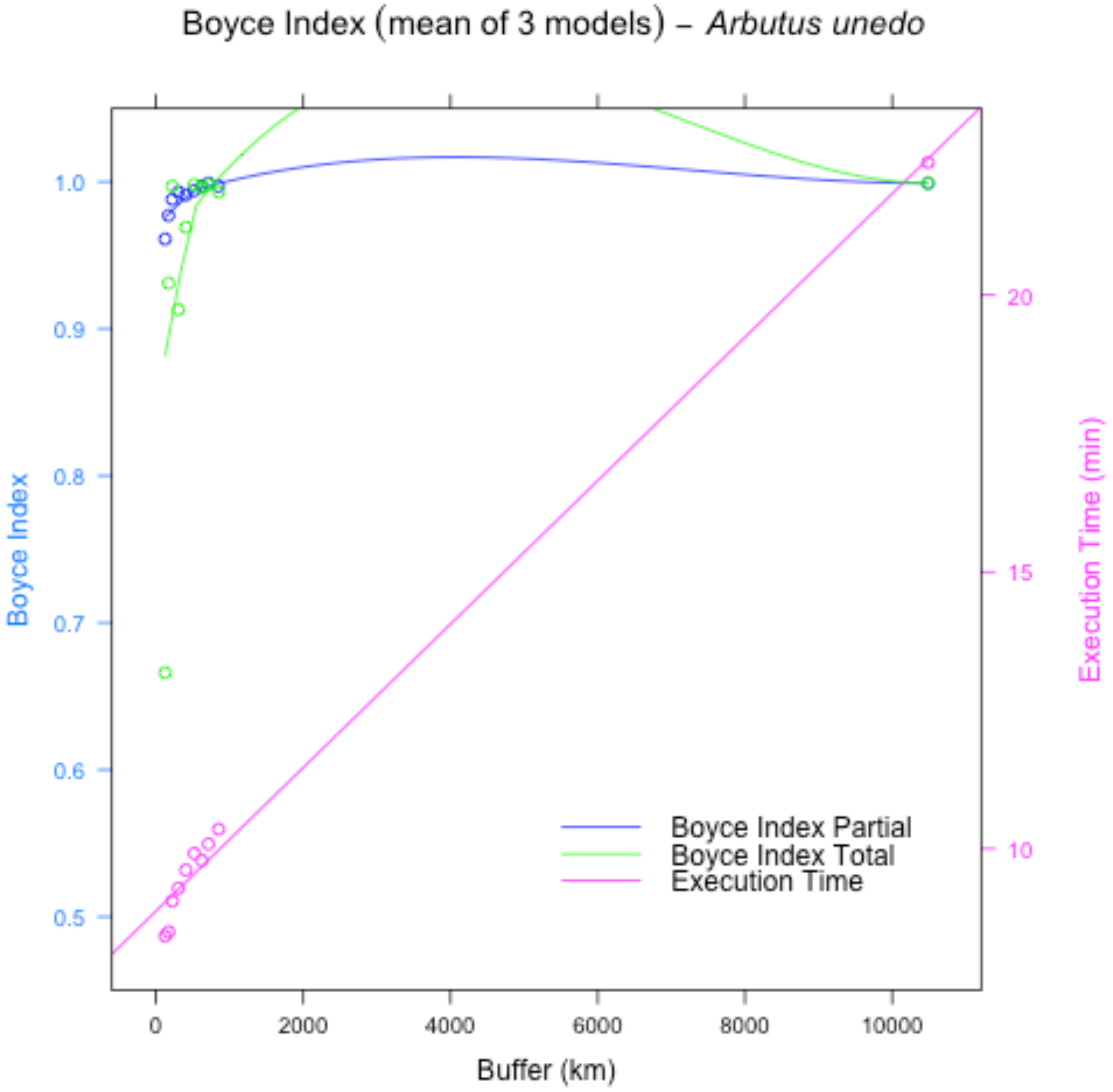

Boyce Index (mean of 3 models) – *Arbutus unedo*
Boyce Index
Execution Time (min)
Buffer (km)
Boyce Index Partial
Boyce Index Total
Execution Time
0.5
0.6
0.7
0.8
0.9
1.0
10
15
20
0
2000
4000
6000
8000
10000


43

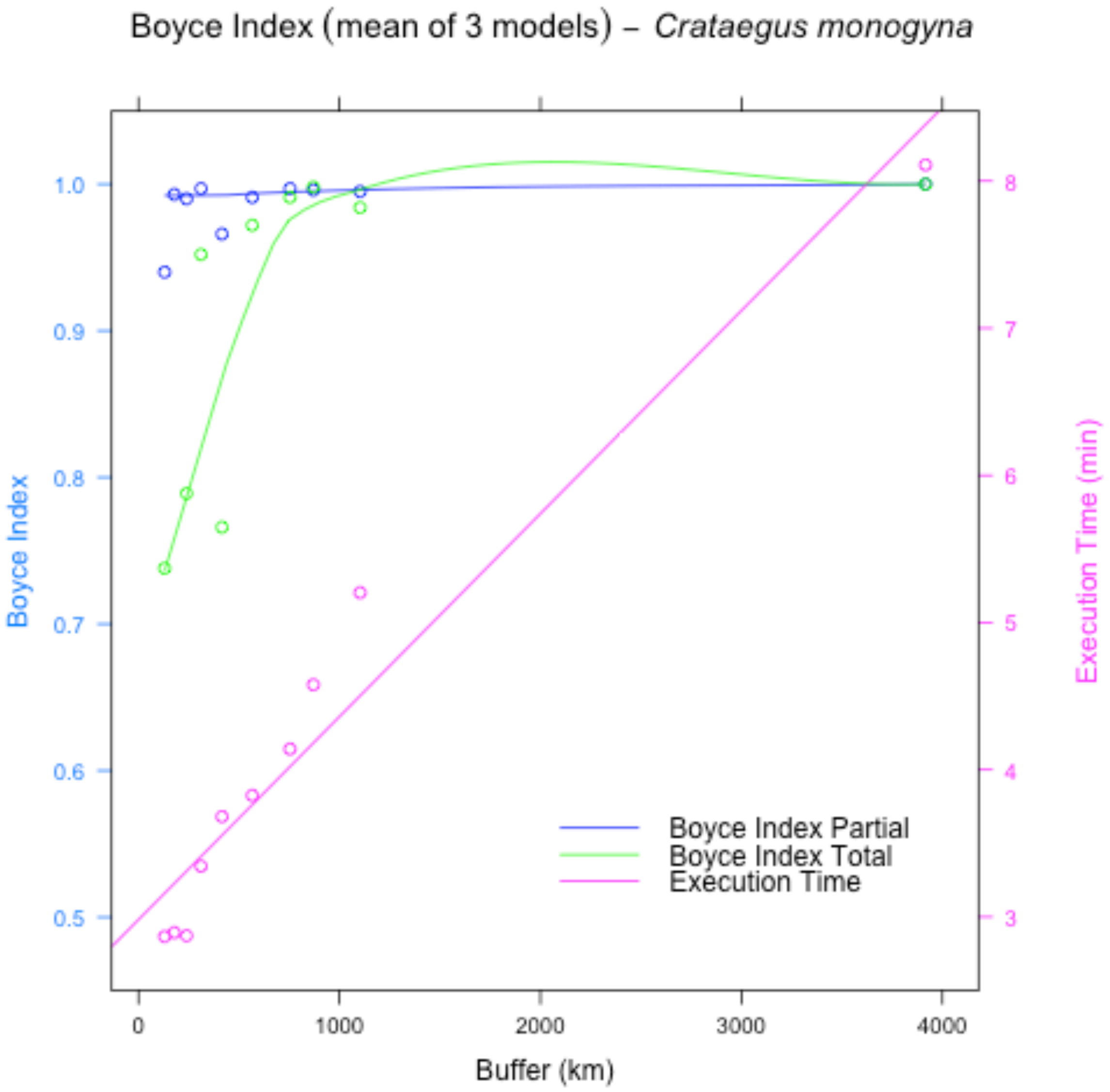
Boyce Index (mean of 3 models) – Crataegus monogyna
Boyce Index
Execution Time (min)
Buffer (km)
Boyce Index Partial
Boyce Index Total
Execution Time
0.5
0.6
0.7
0.8
0.9
1.0
3
4
5
6
7
8
0
1000
2000
3000
4000

44

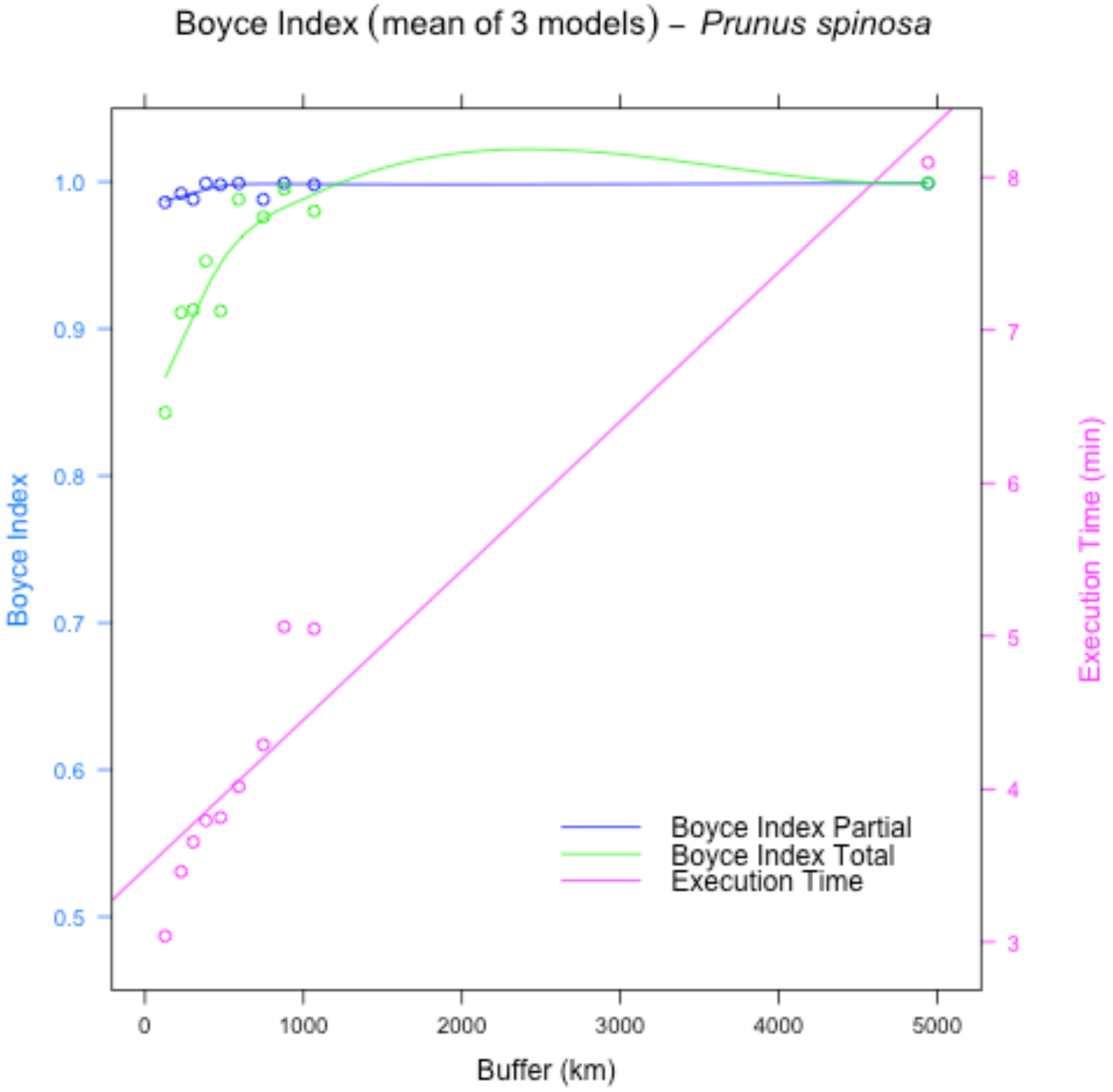
Boyce Index (mean of 3 models) – *Prunus spinosa*
Boyce Index
Execution Time (min)
Buffer (km)
Boyce Index Partial
Boyce Index Total
Execution Time
0
1000
2000
3000
4000
5000
0.5
0.6
0.7
0.8
0.9
1.0
3
4
5
6
7
8

45

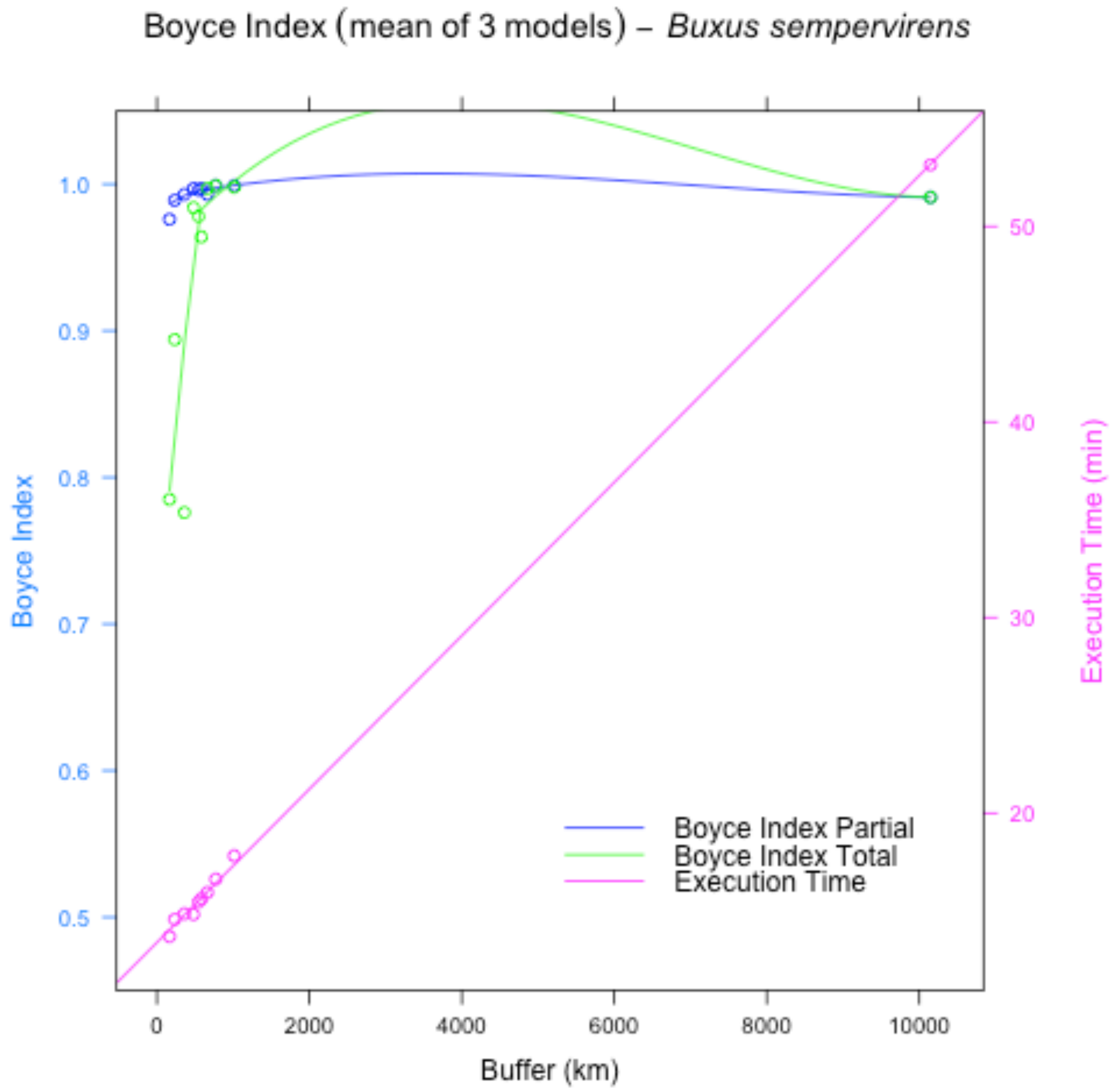
Boyce Index (mean of 3 models) – *Buxus sempervirens*
Boyce Index
1.0
0.9
0.8
0.7
0.6
0.5
Execution Time (min)
50
40
30
20
0
2000
4000
6000
8000
10000
Buffer (km)
Boyce Index Partial
Boyce Index Total
Execution Time

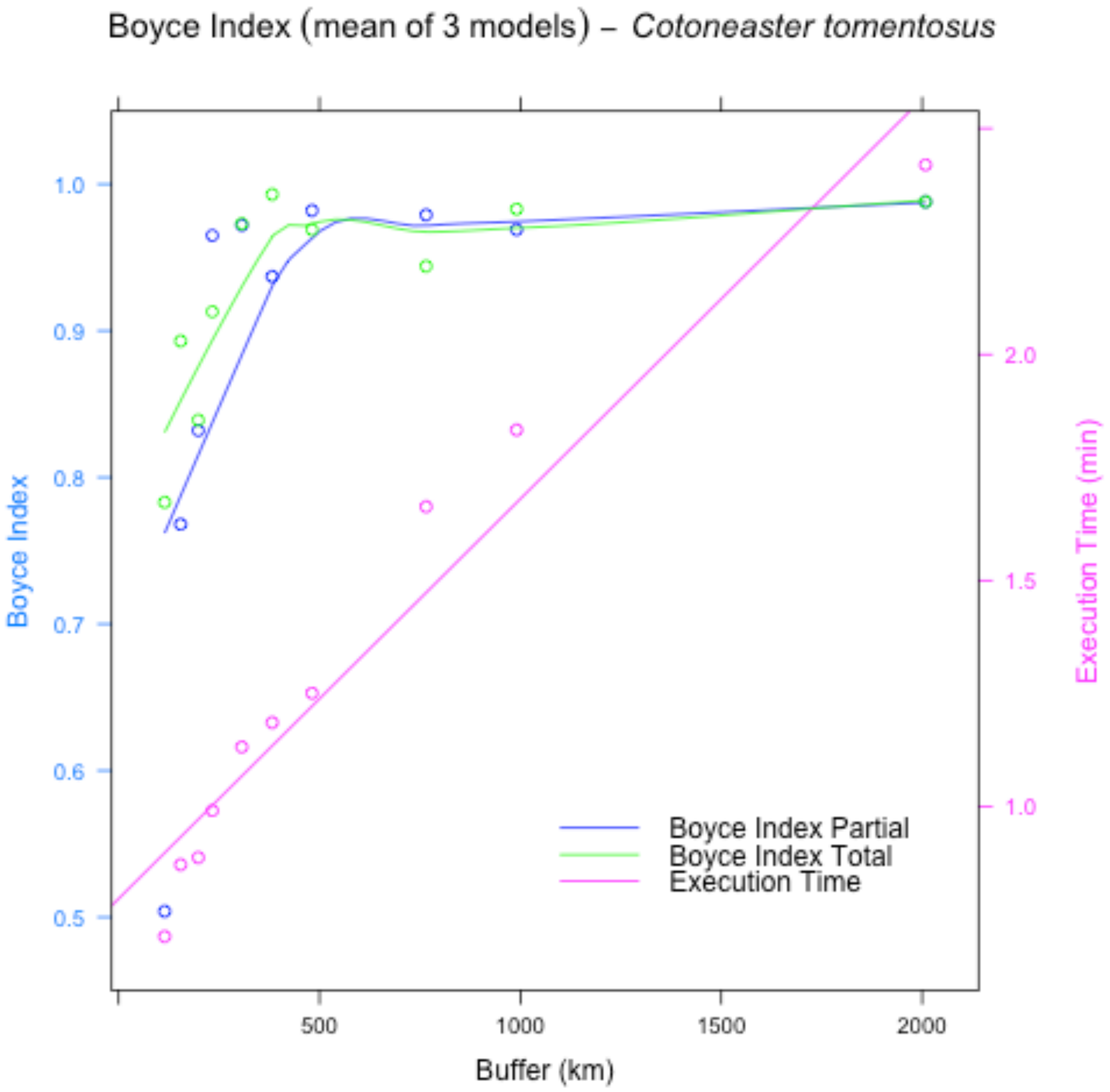

Boyce Index (mean of 3 models) – Cotoneaster tomentosus
Boyce Index
1.0
0.9
0.8
0.7
0.6
0.5
Execution Time (min)
2.0
1.5
1.0
500
1000
1500
2000
Buffer (km)
Boyce Index Partial
Boyce Index Total
Execution Time


47

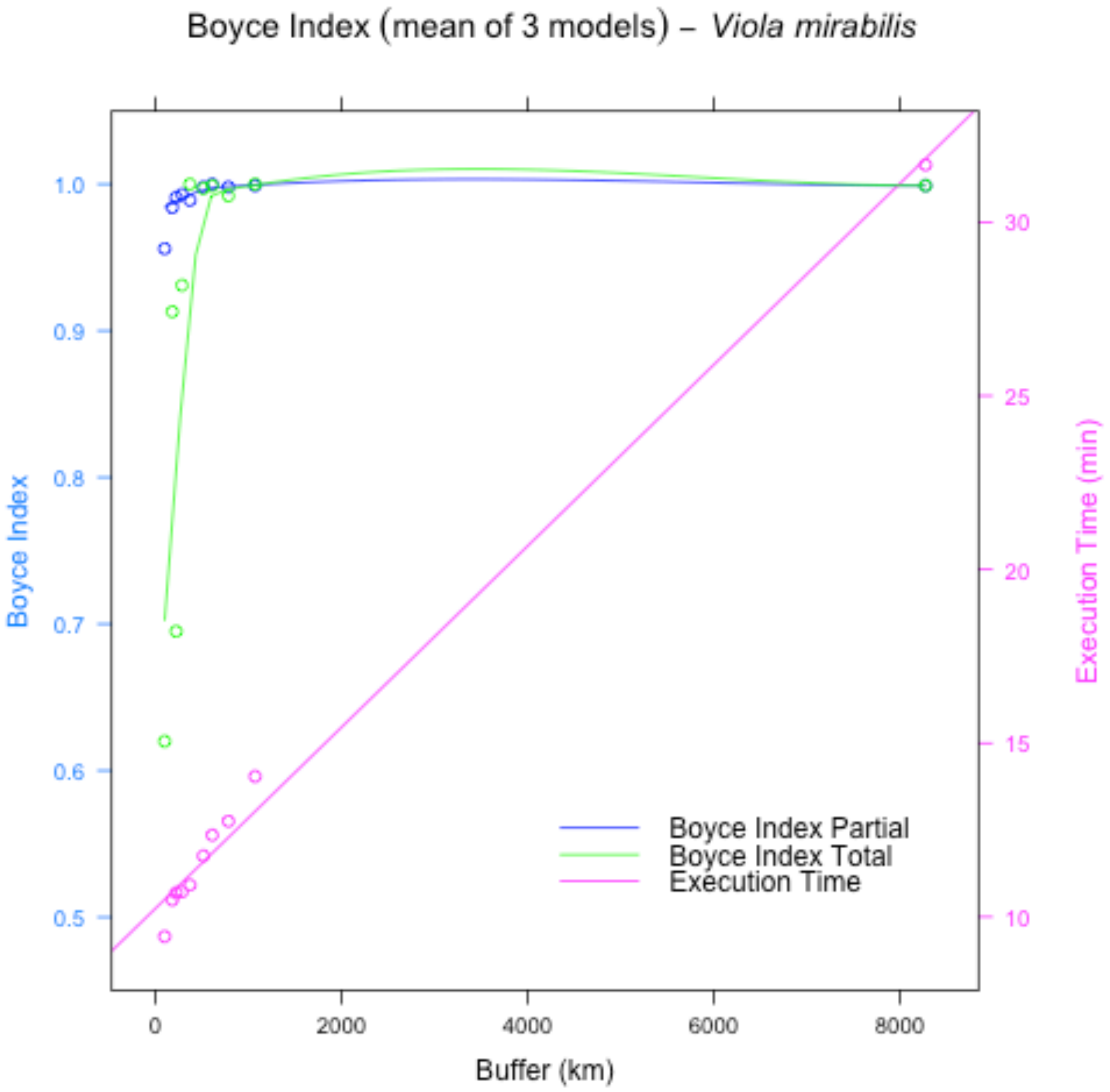

Boyce Index (mean of 3 models) – *Viola mirabilis*
Boyce Index
Execution Time (min)
Buffer (km)
Boyce Index Partial
Boyce Index Total
Execution Time
0
2000
4000
6000
8000
0.5
0.6
0.7
0.8
0.9
1.0
10
15
20
25
30


48

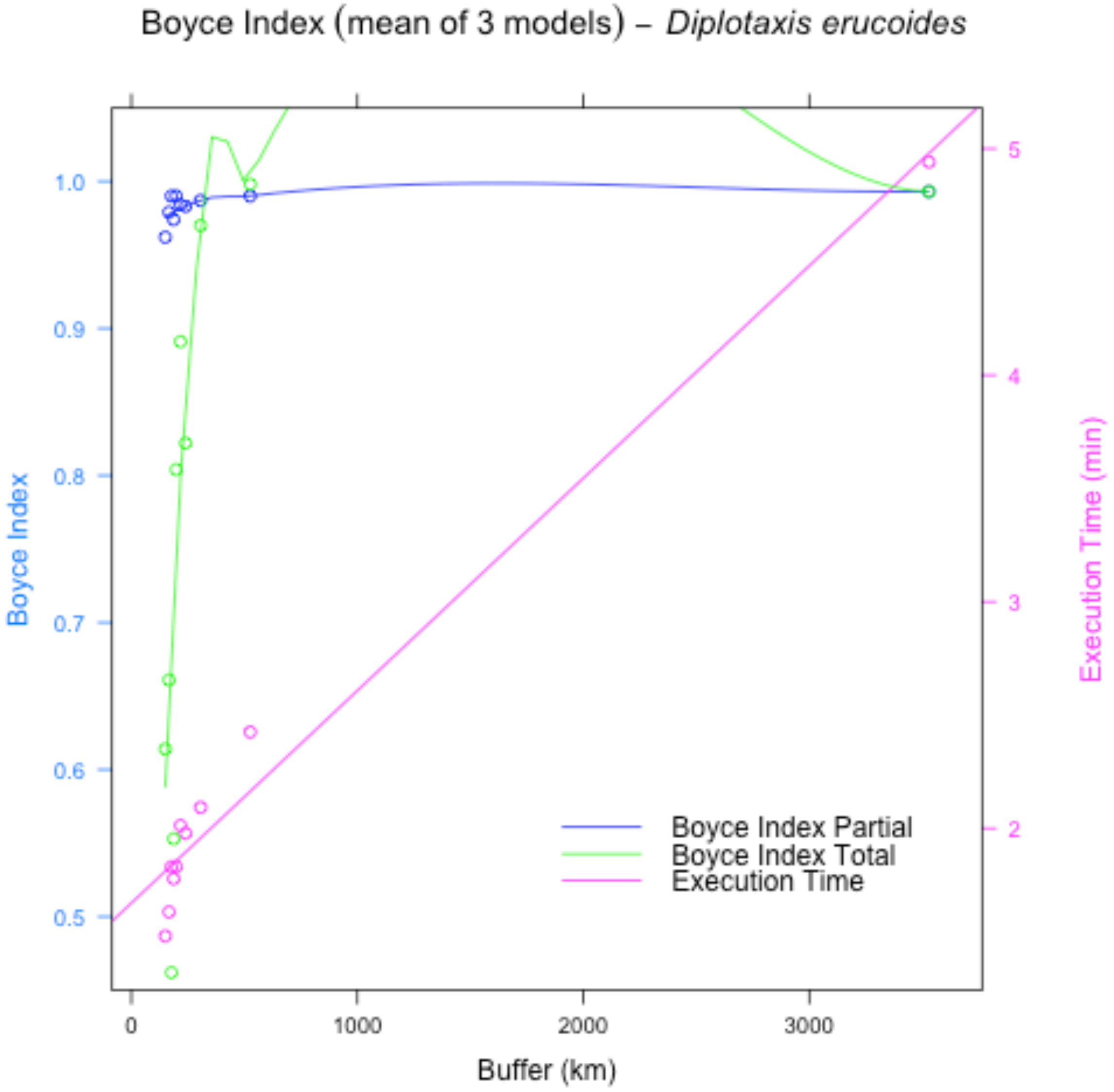

Boyce Index (mean of 3 models) – *Diplotaxis erucoides*
Boyce Index
1.0
0.9
0.8
0.7
0.6
0.5
Execution Time (min)
5
4
3
2
0
1000
2000
3000
Buffer (km)
Boyce Index Partial
Boyce Index Total
Execution Time


49

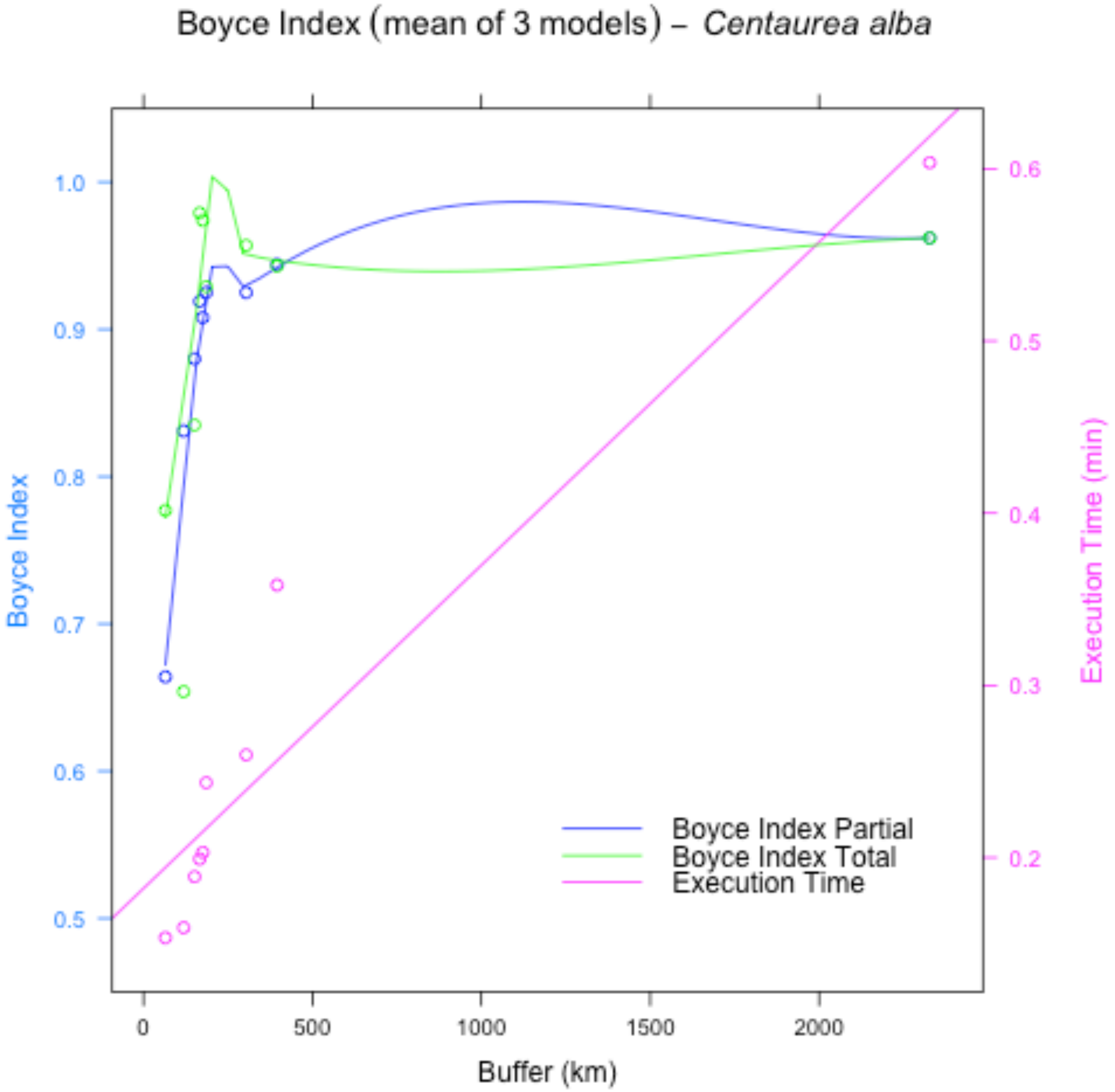
Boyce Index (mean of 3 models) – Centaurea alba
Boyce Index
Execution Time (min)
Buffer (km)
1.0
0.9
0.8
0.7
0.6
0.5
0.6
0.5
0.4
0.3
0.2
0
500
1000
1500
2000
Boyce Index Partial
Boyce Index Total
Execution Time

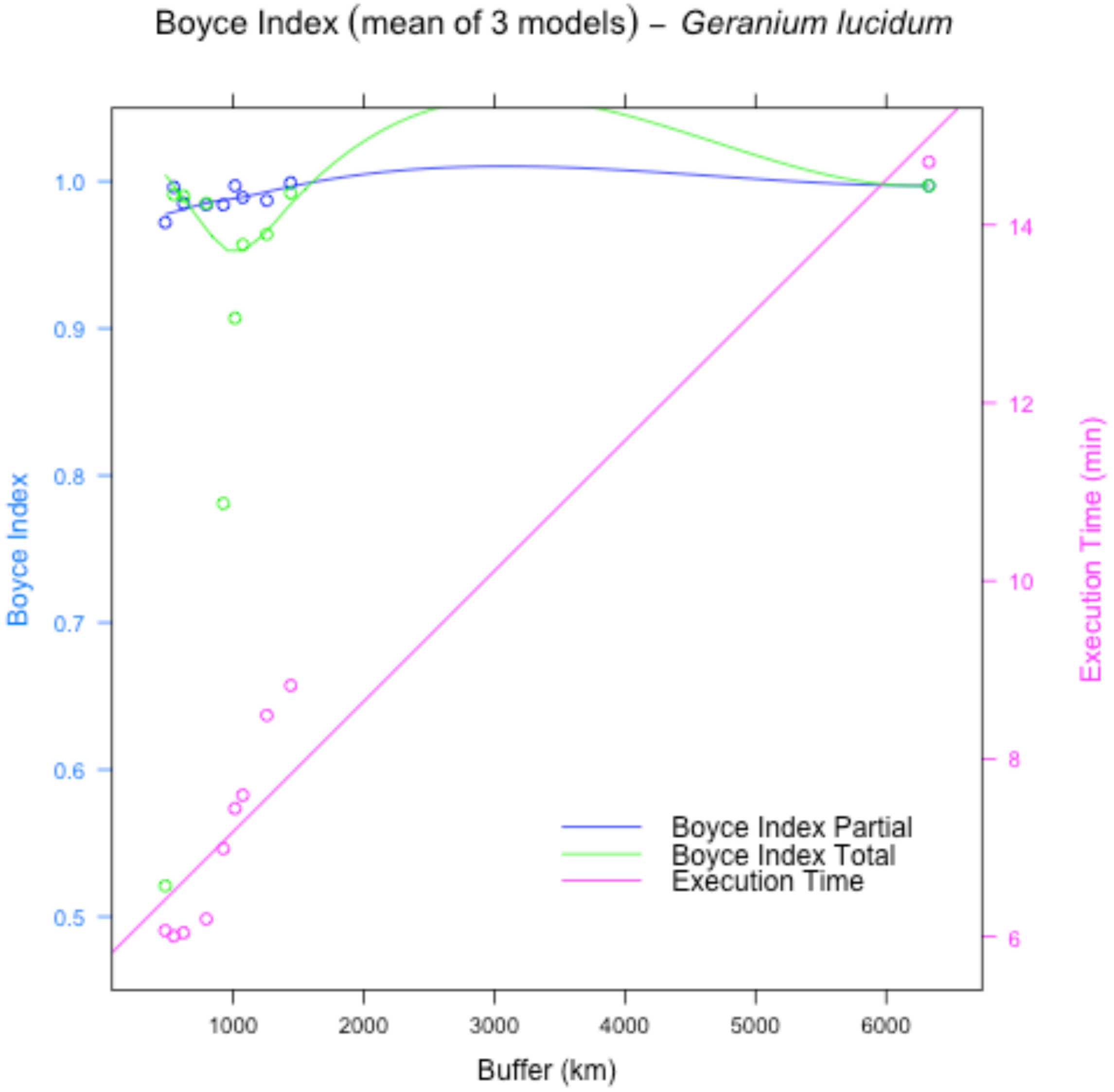

Boyce Index (mean of 3 models) – Geranium lucidum
Boyce Index
1.0
0.9
0.8
0.7
0.6
0.5
Execution Time (min)
14
12
10
8
6
1000
2000
3000
4000
5000
6000
Buffer (km)
Boyce Index Partial
Boyce Index Total
Execution Time

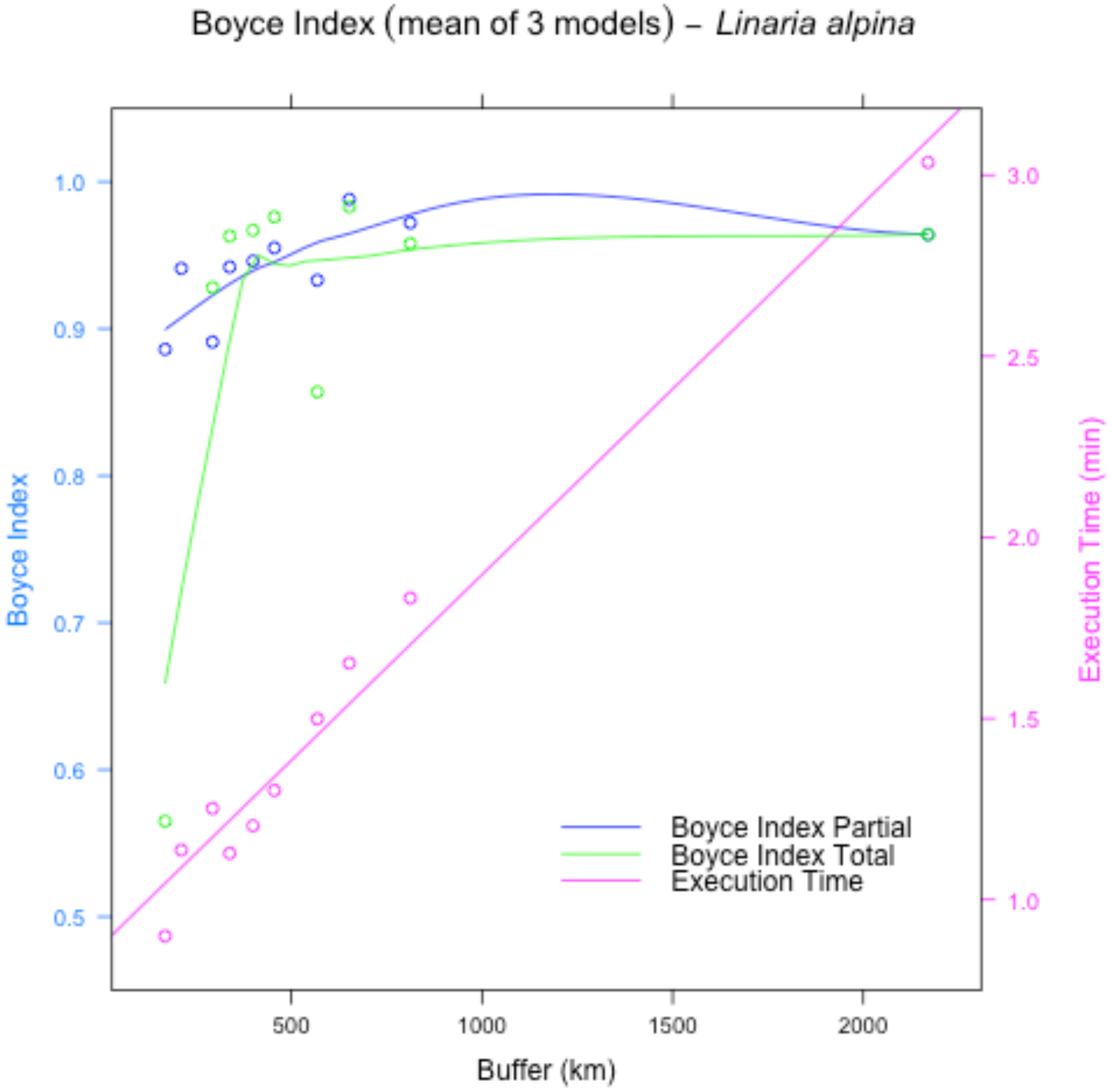
Boyce Index (mean of 3 models) – Linaria alpina
Boyce Index
Execution Time (min)
Buffer (km)
Boyce Index Partial
Boyce Index Total
Execution Time
0.5
0.6
0.7
0.8
0.9
1.0
1.0
1.5
2.0
2.5
3.0
500
1000
1500
2000

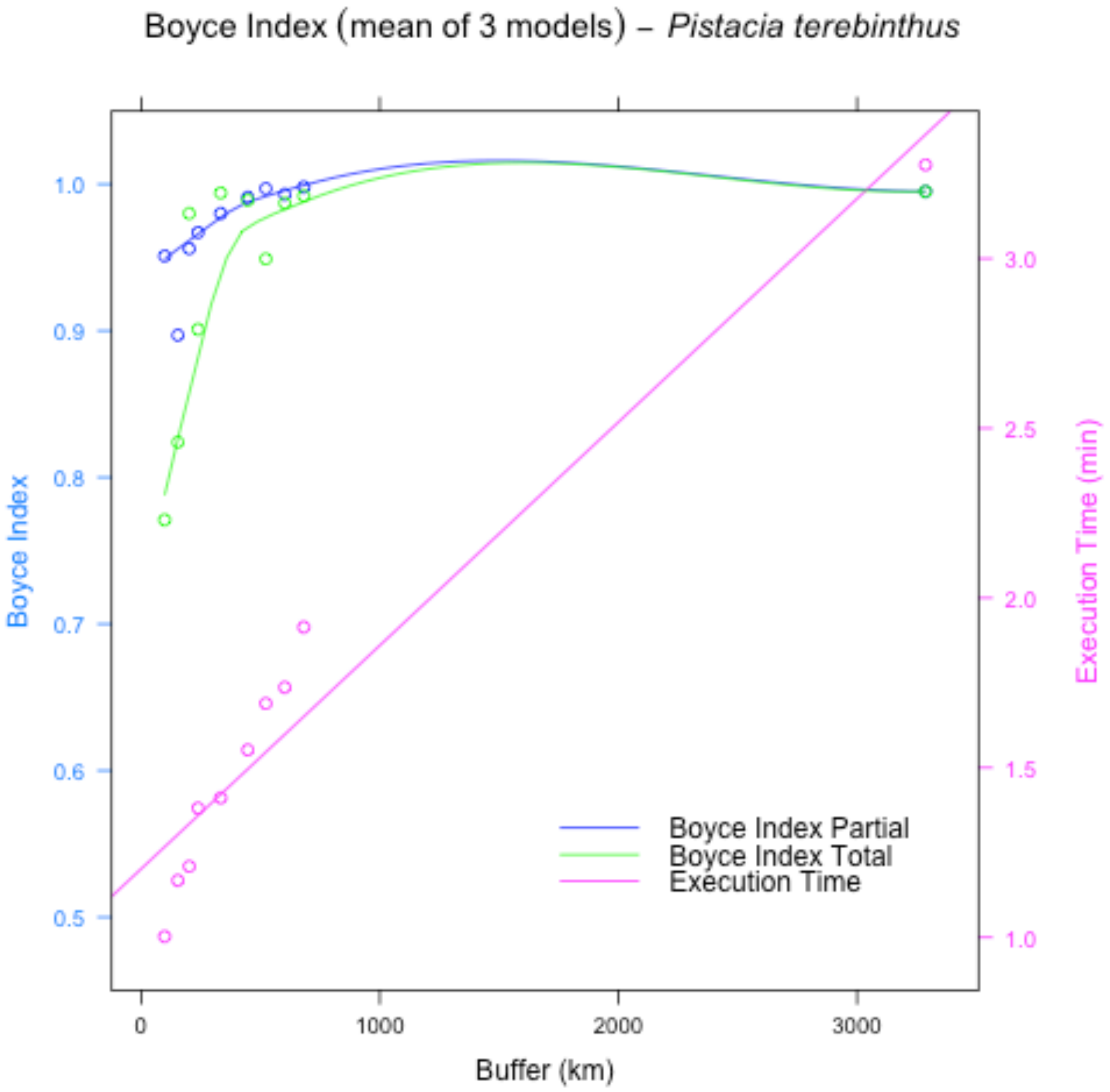
Boyce Index (mean of 3 models) – Pistacia terebinthus
Boyce Index
Execution Time (min)
Buffer (km)
Boyce Index Partial
Boyce Index Total
Execution Time
1.0
0.9
0.8
0.7
0.6
0.5
3.0
2.5
2.0
1.5
1.0
0
1000
2000
3000

53

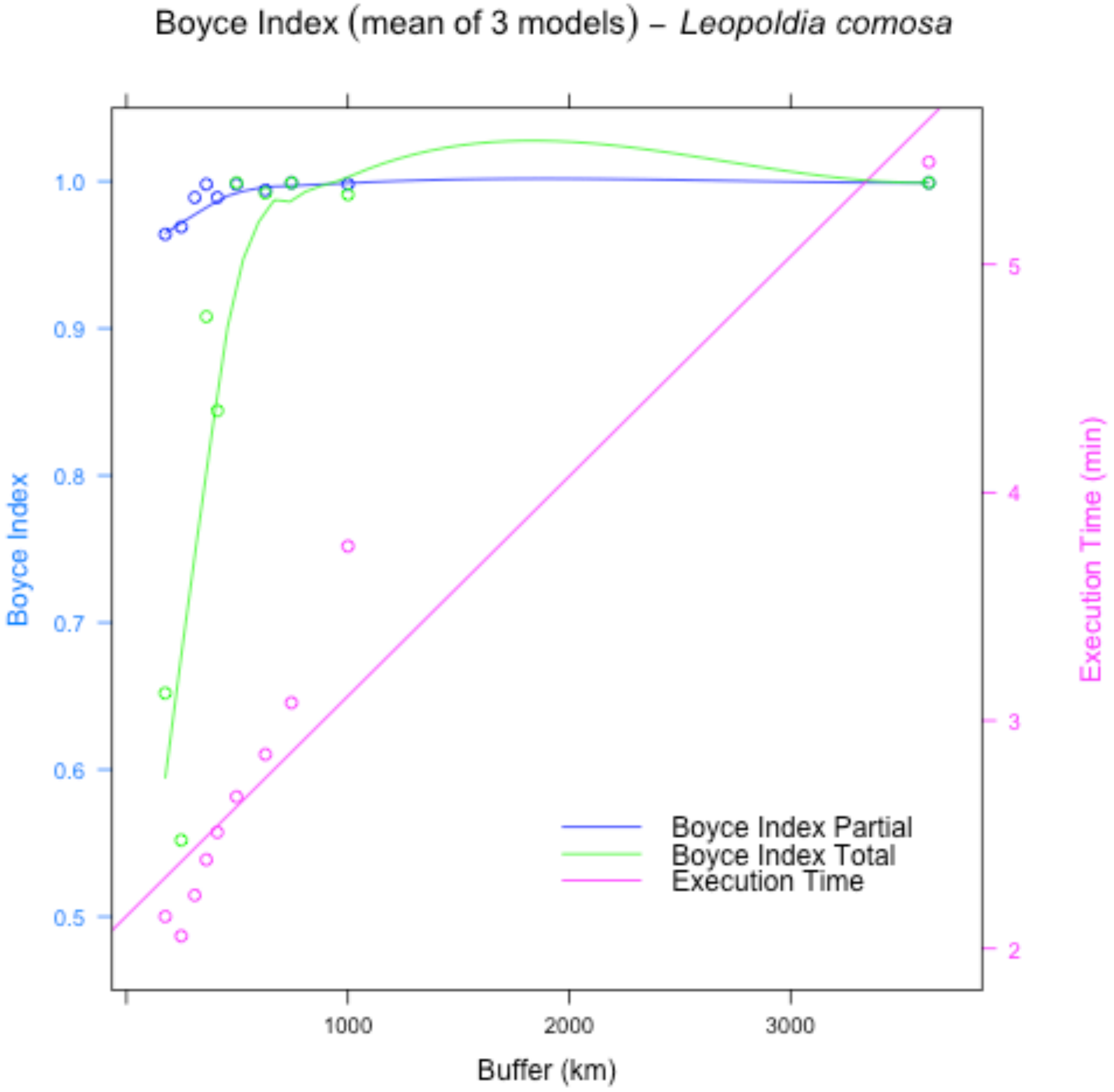

Boyce Index (mean of 3 models) – *Leopoldia comosa*
1.0
0.9
0.8
0.7
0.6
0.5
Boyce Index
5
4
3
2
Execution Time (min)
Boyce Index Partial
Boyce Index Total
Execution Time
1000
2000
3000
Buffer (km)


54

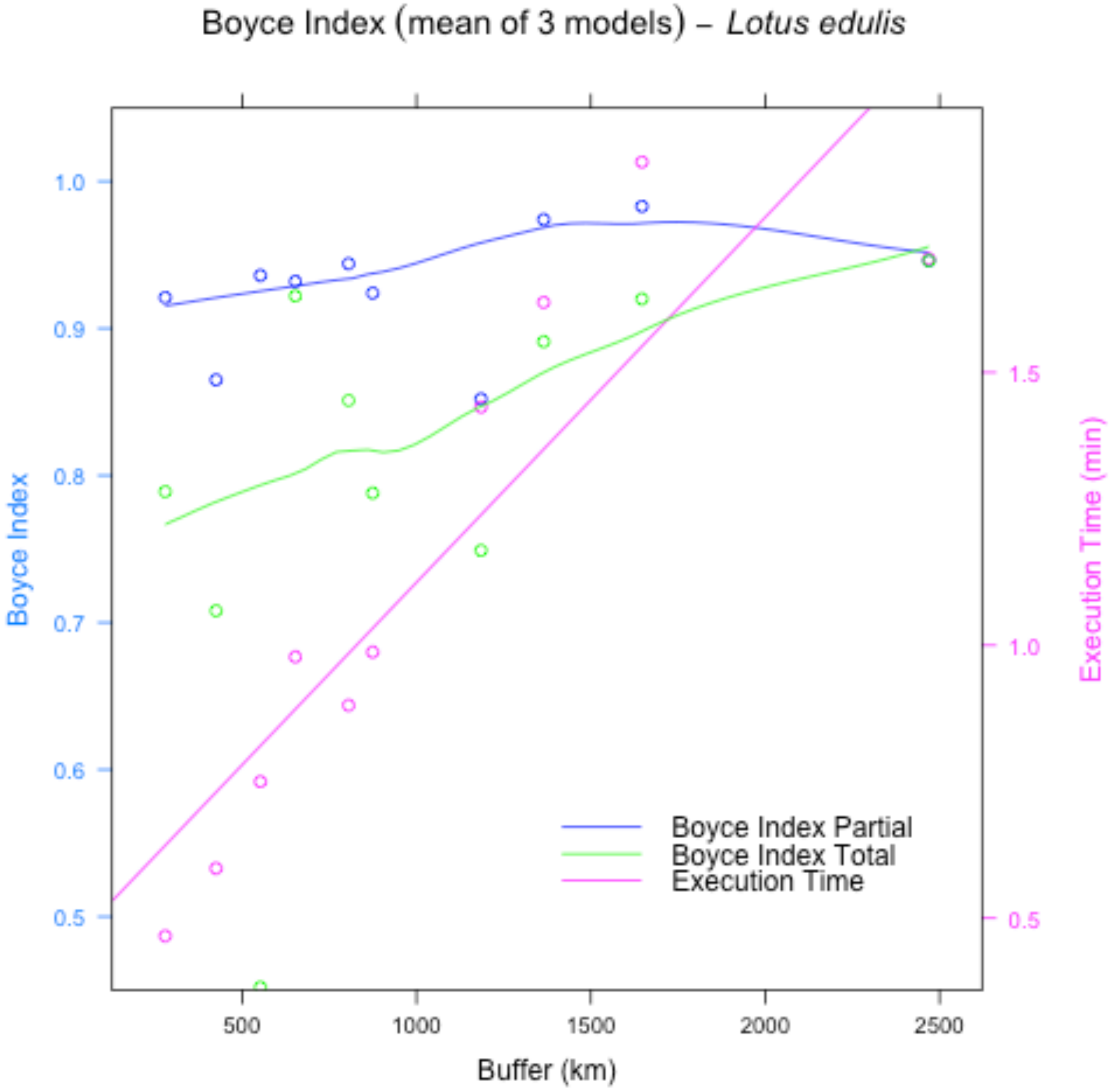
Boyce Index (mean of 3 models) – Lotus edulis
Boyce Index
Execution Time (min)
Buffer (km)
1.0
0.9
0.8
0.7
0.6
0.5
1.5
1.0
0.5
500
1000
1500
2000
2500
Boyce Index Partial
Boyce Index Total
Execution Time

57 # Supplementary Material S4

58 Figures S4.1 - S4.10: Evolution of Boyce Index Total (green) and Parcial (blue), and the
59 execution time in minutes (pink), for all the species in Case Study 2

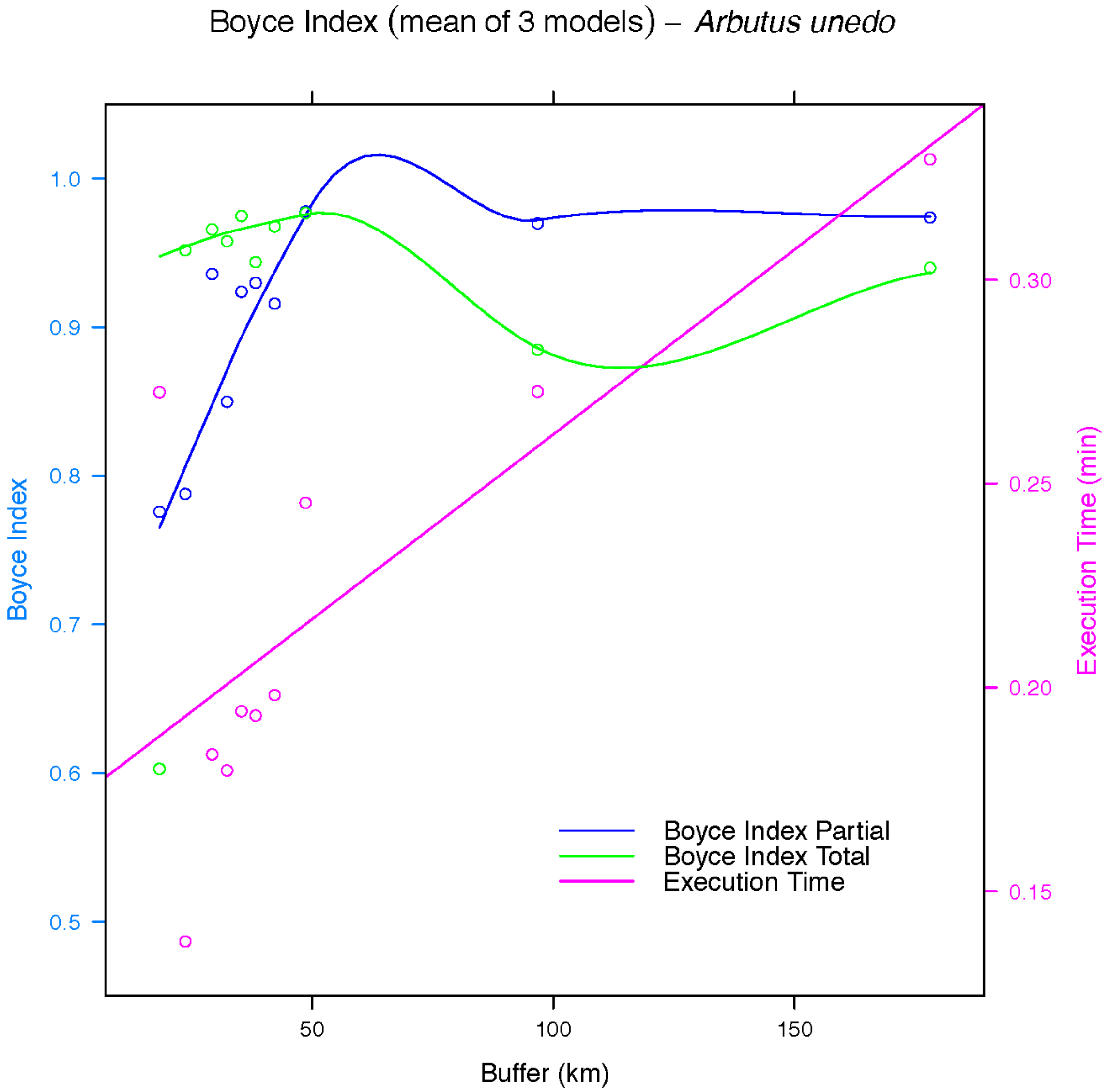


60

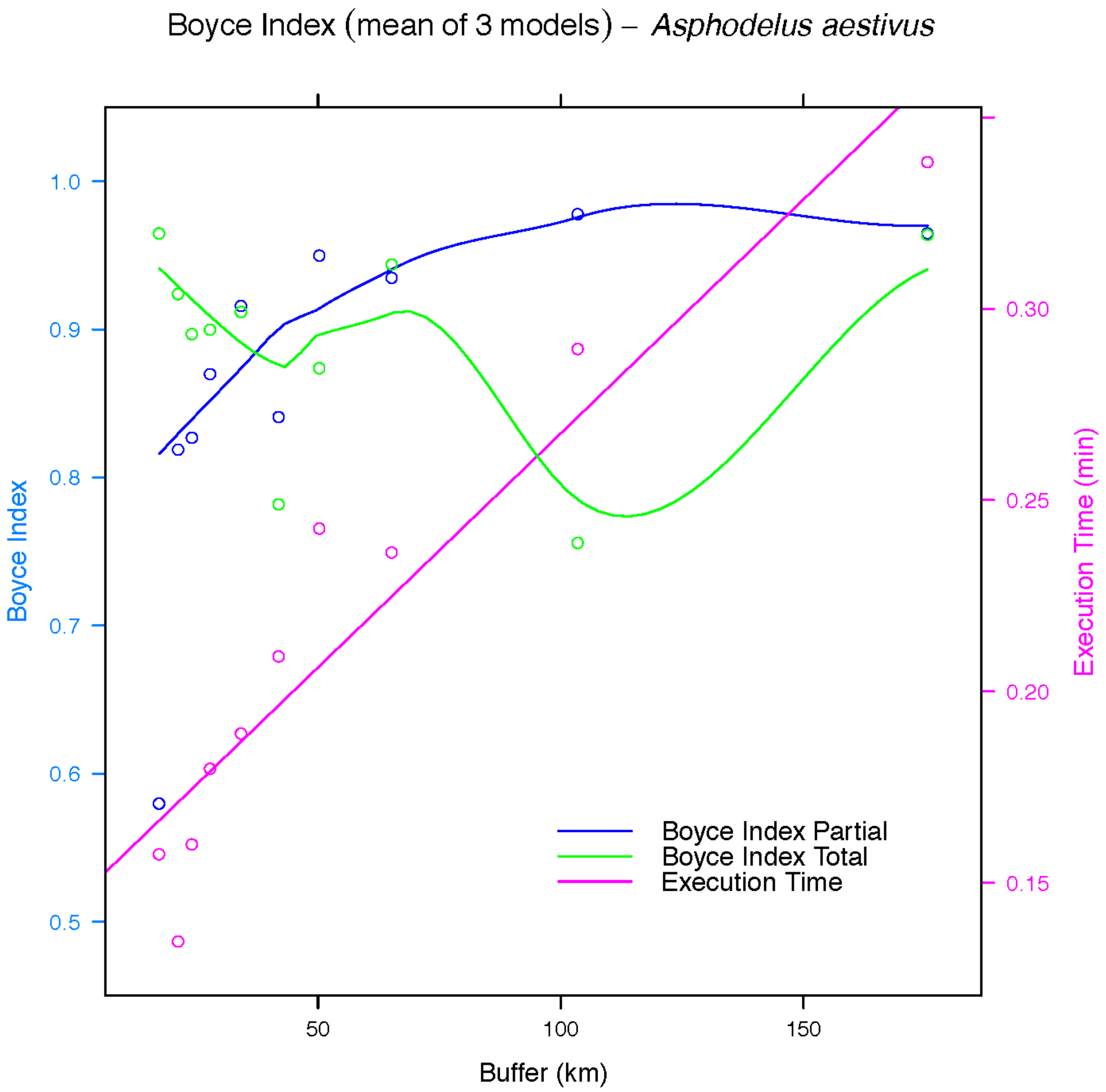
Boyce Index (mean of 3 models) – *Asphodelus aestivus*
Boyce Index
1.0
0.9
0.8
0.7
0.6
0.5
Execution Time (min)
0.30
0.25
0.20
0.15
50
100
150
Buffer (km)
Boyce Index Partial
Boyce Index Total
Execution Time

61

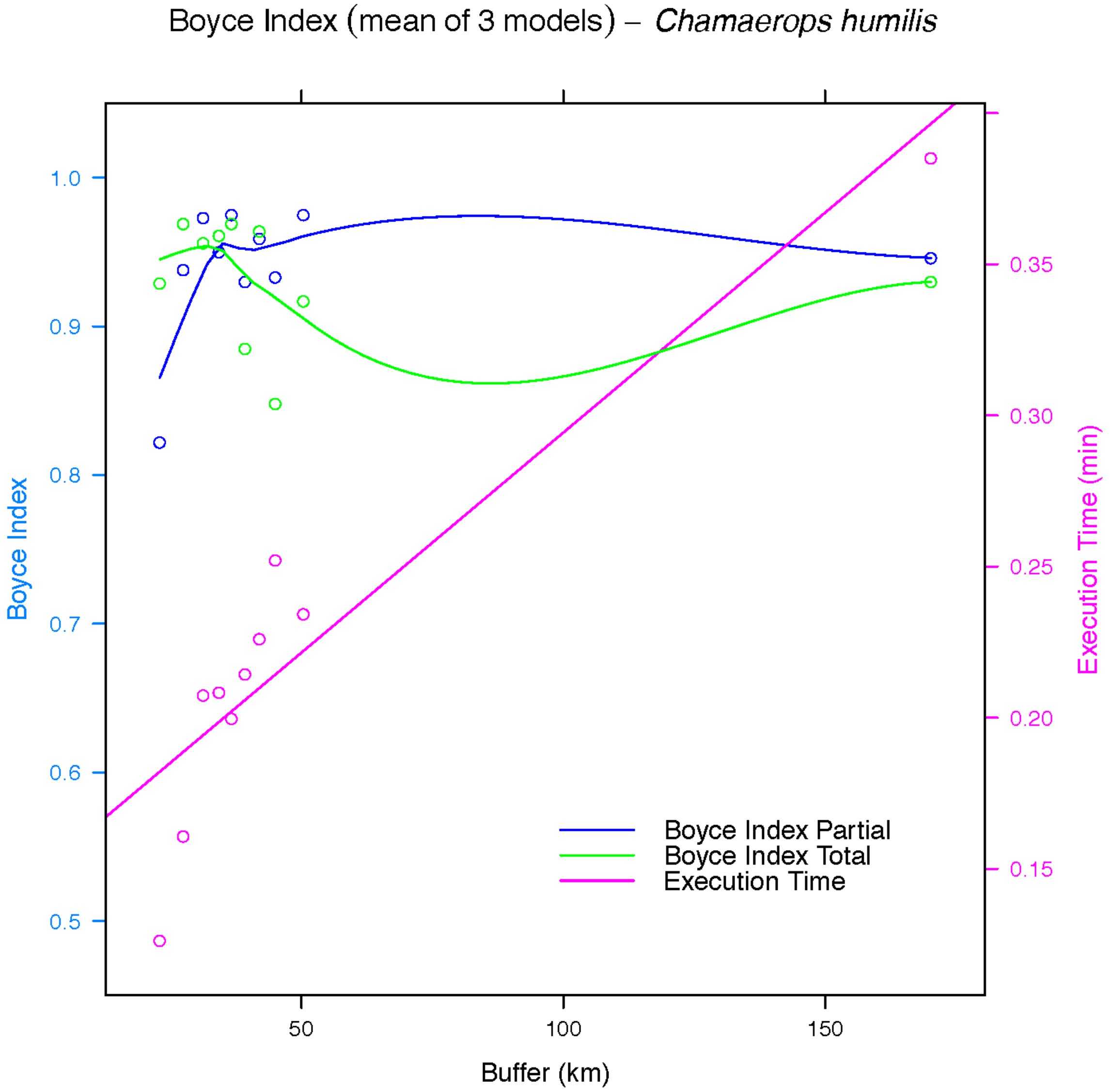

Boyce Index (mean of 3 models) – Chamaerops humilis
Boyce Index
1.0
0.9
0.8
0.7
0.6
0.5
Execution Time (min)
0.35
0.30
0.25
0.20
0.15
50
100
150
Buffer (km)
Boyce Index Partial
Boyce Index Total
Execution Time


62

Boyce Index (mean of 3 models) – *Ephedra fragilis subsp. fragilis*

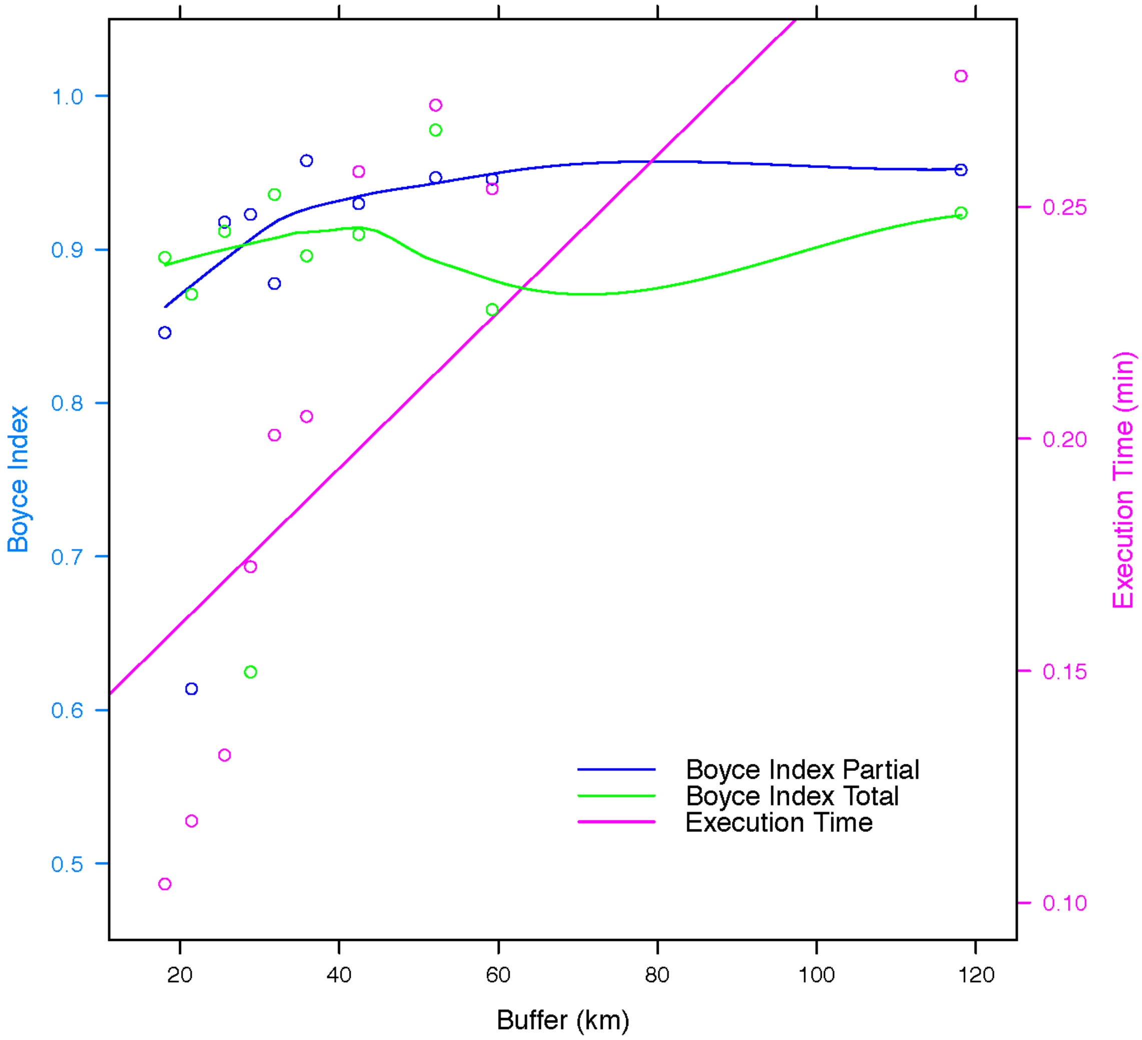


63

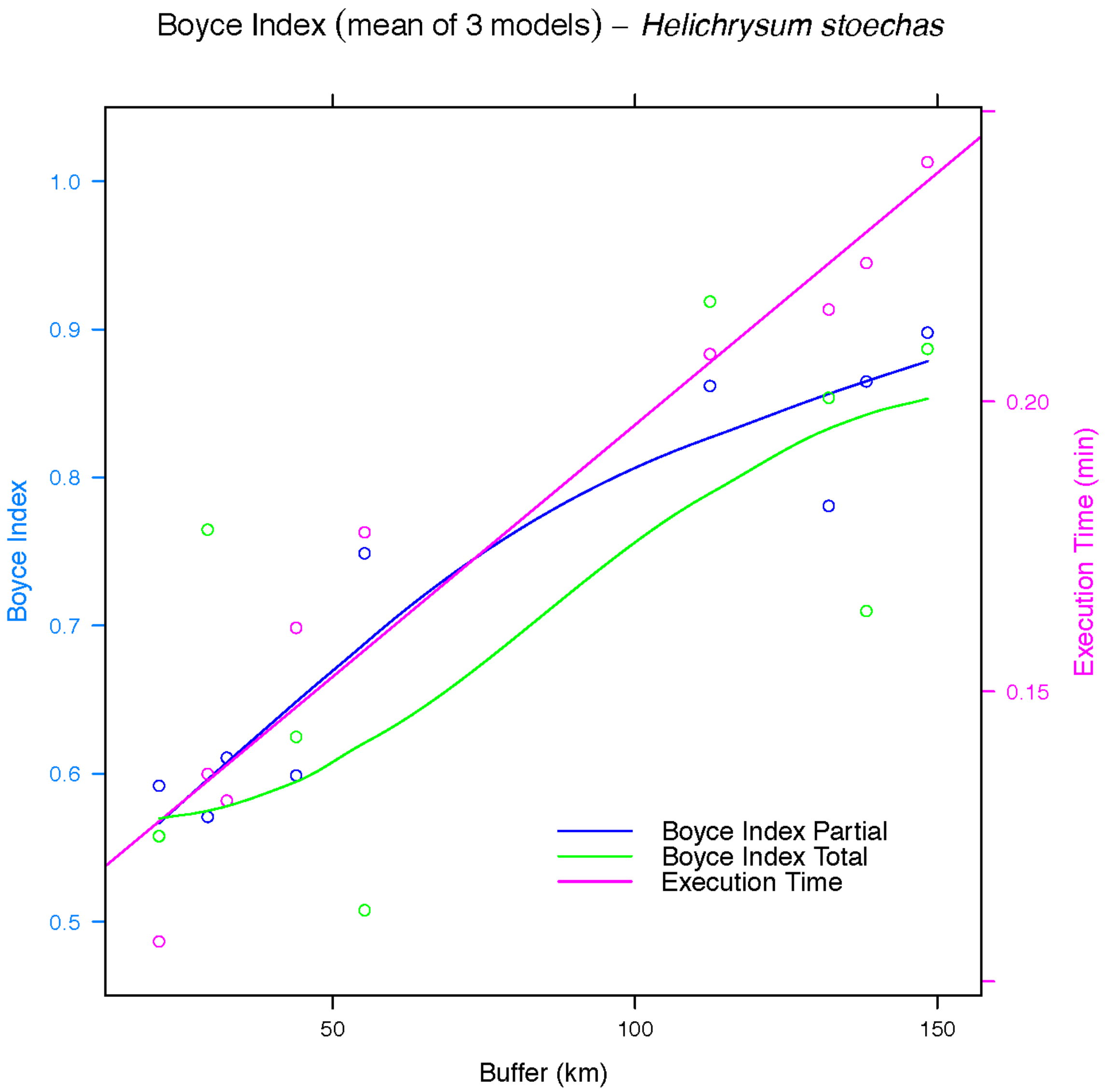

Boyce Index (mean of 3 models) – *Helichrysum stoechas*
Boyce Index
1.0
0.9
0.8
0.7
0.6
0.5
Execution Time (min)
0.20
0.15
50
100
150
Buffer (km)
Boyce Index Partial
Boyce Index Total
Execution Time


64

Boyce Index (mean of 3 models) – *Juniperus oxycedrus subsp. oxycedrus*

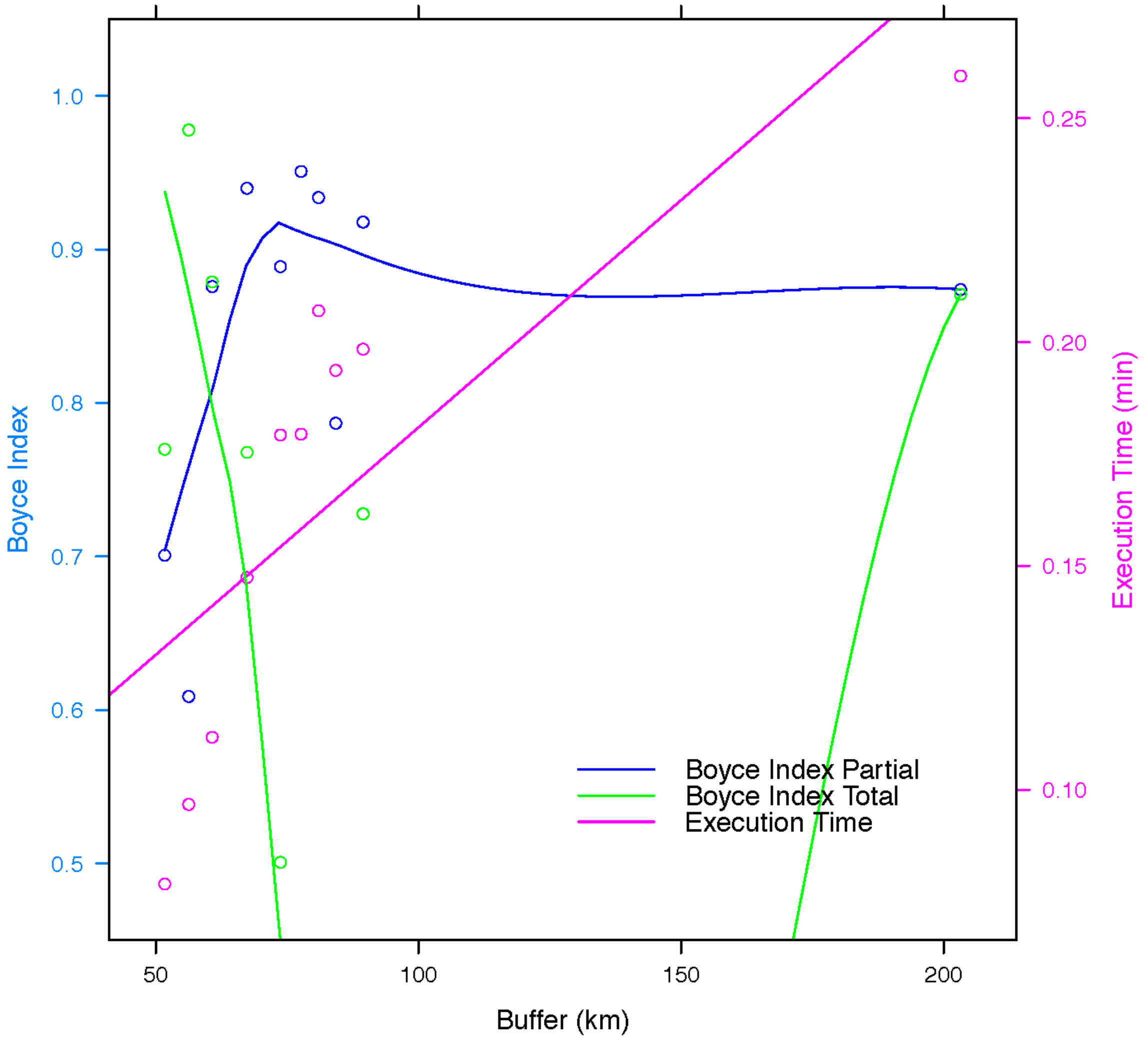


65

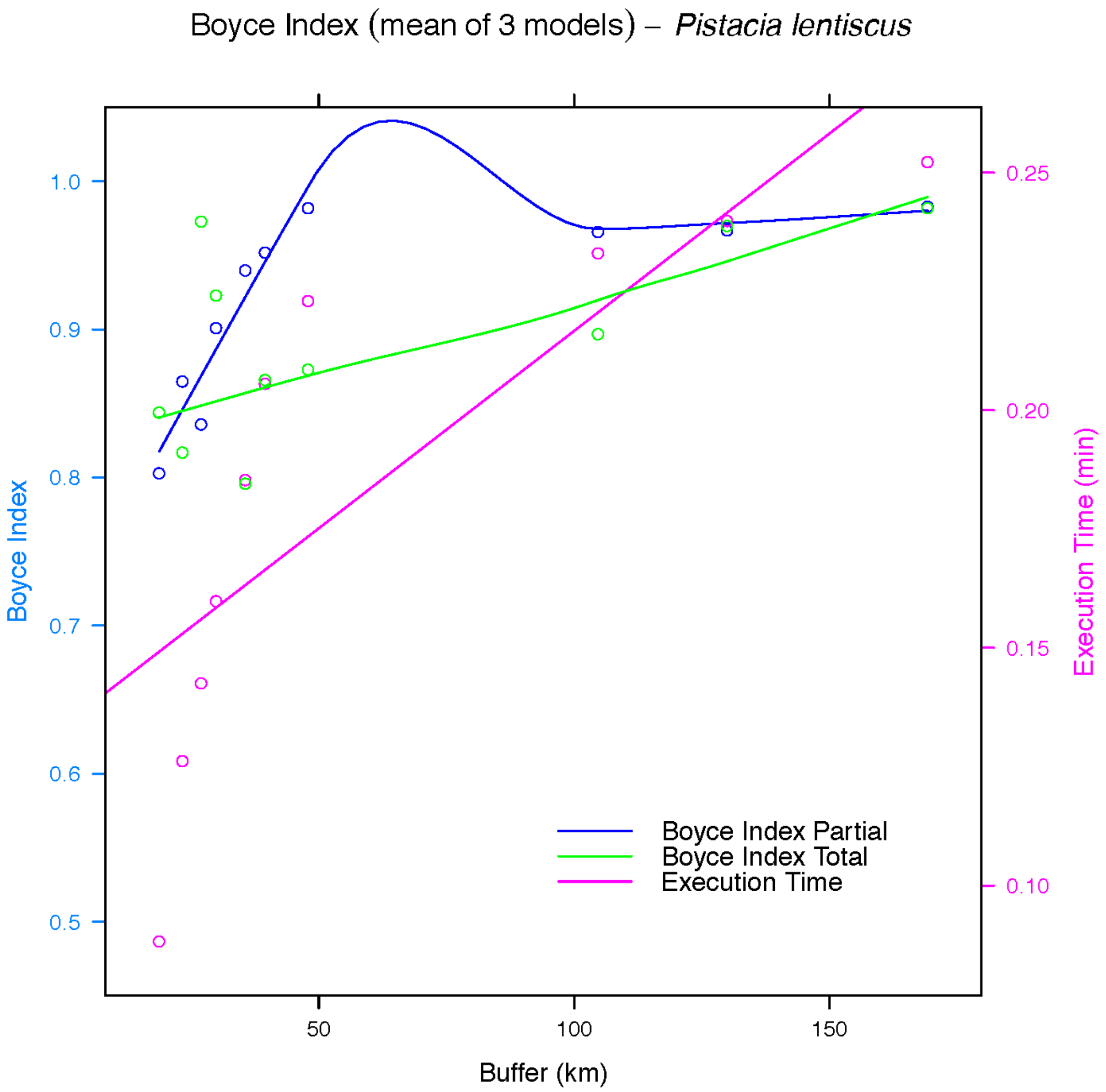
Boyce Index (mean of 3 models) – *Pistacia lentiscus*
Boyce Index
1.0
0.9
0.8
0.7
0.6
0.5
Execution Time (min)
0.25
0.20
0.15
0.10
50
100
150
Buffer (km)
Boyce Index Partial
Boyce Index Total
Execution Time

66

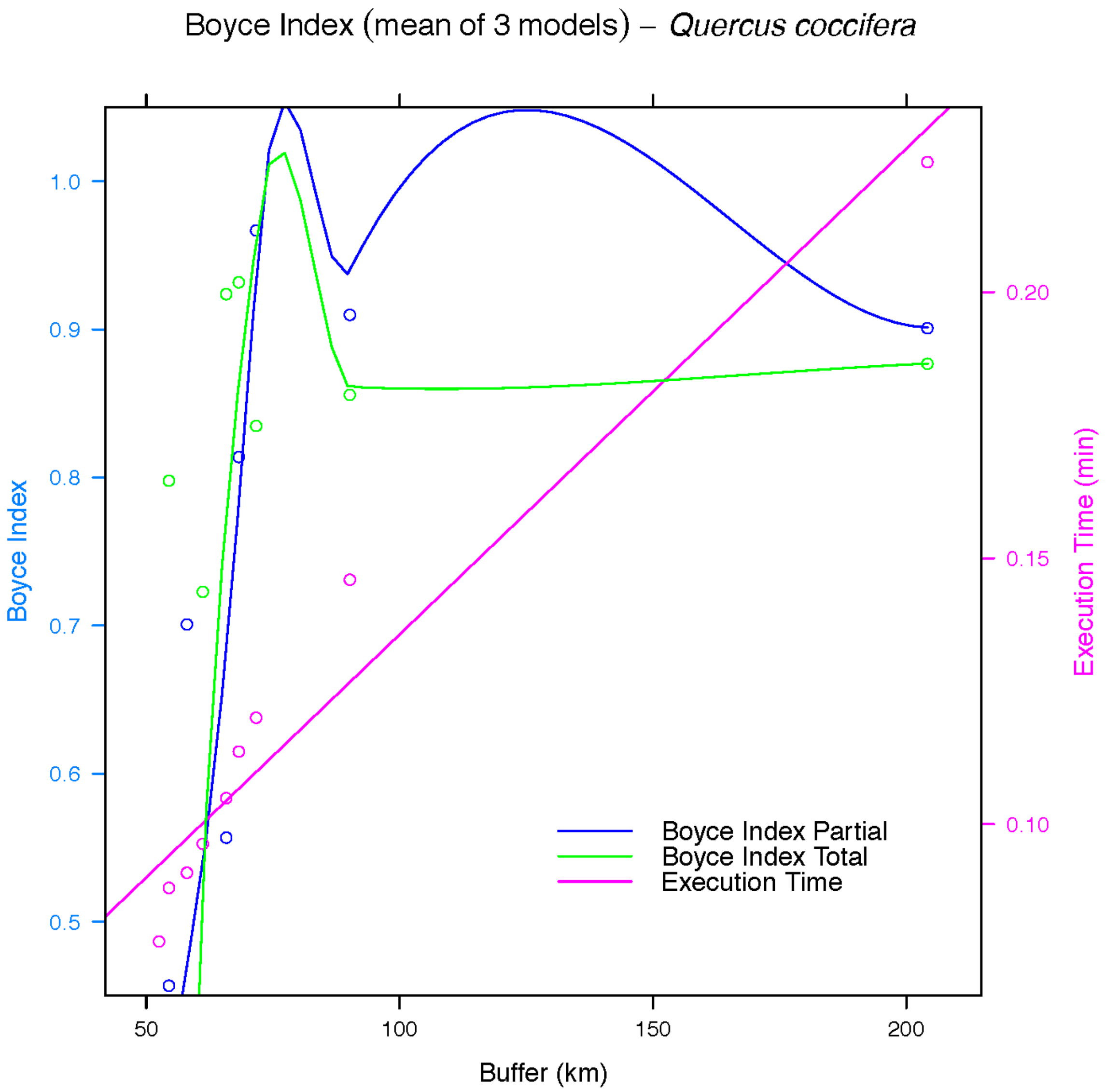
Boyce Index (mean of 3 models) – *Quercus coccifera*
Boyce Index
1.0
0.9
0.8
0.7
0.6
0.5
Execution Time (min)
0.20
0.15
0.10
50
100
150
200
Buffer (km)
Boyce Index Partial
Boyce Index Total
Execution Time

67

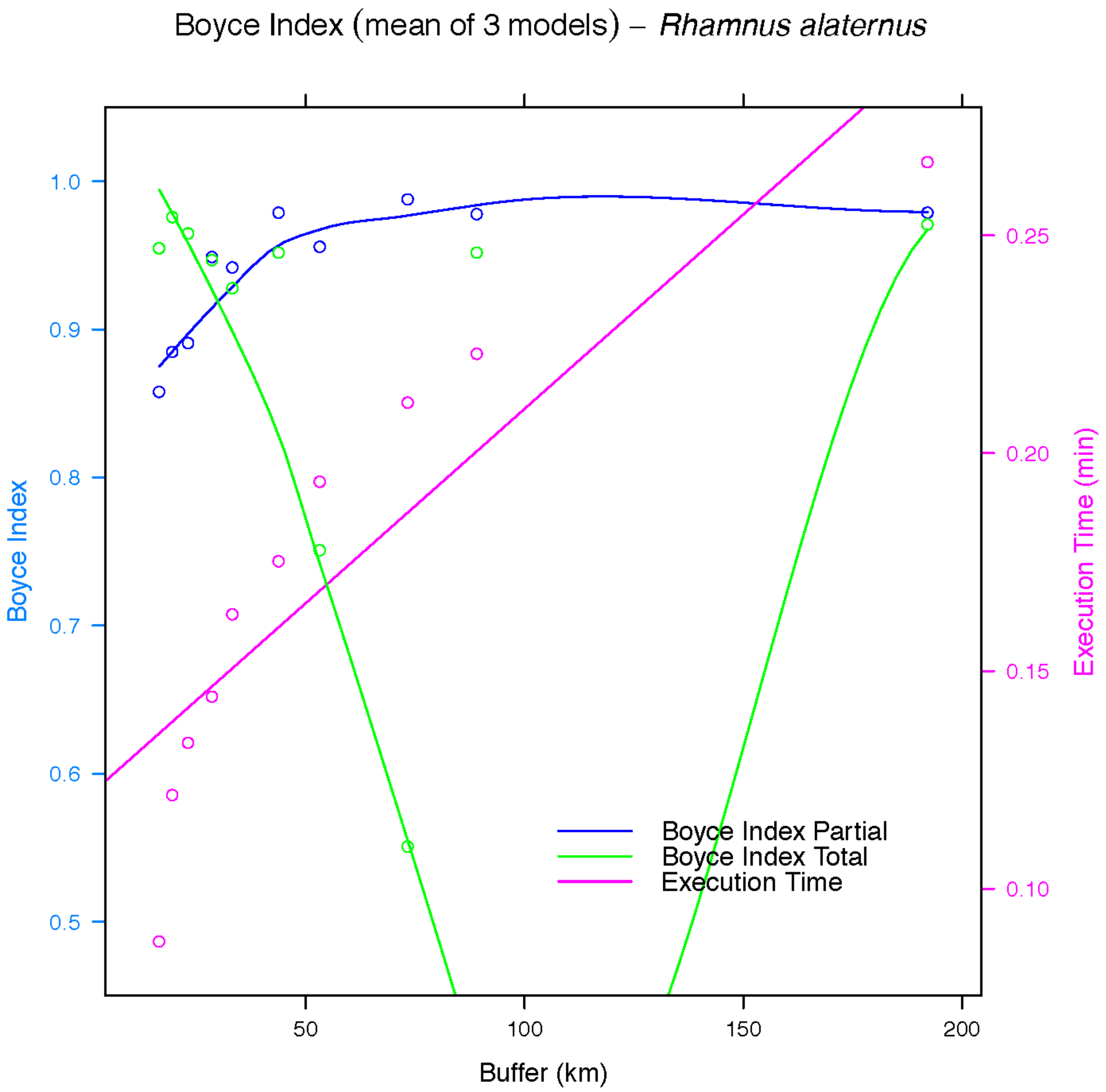

Boyce Index (mean of 3 models) – *Rhamnus alaternus*
Boyce Index
1.0
0.9
0.8
0.7
0.6
0.5
Execution Time (min)
0.25
0.20
0.15
0.10
50
100
150
200
Buffer (km)
Boyce Index Partial
Boyce Index Total
Execution Time


68

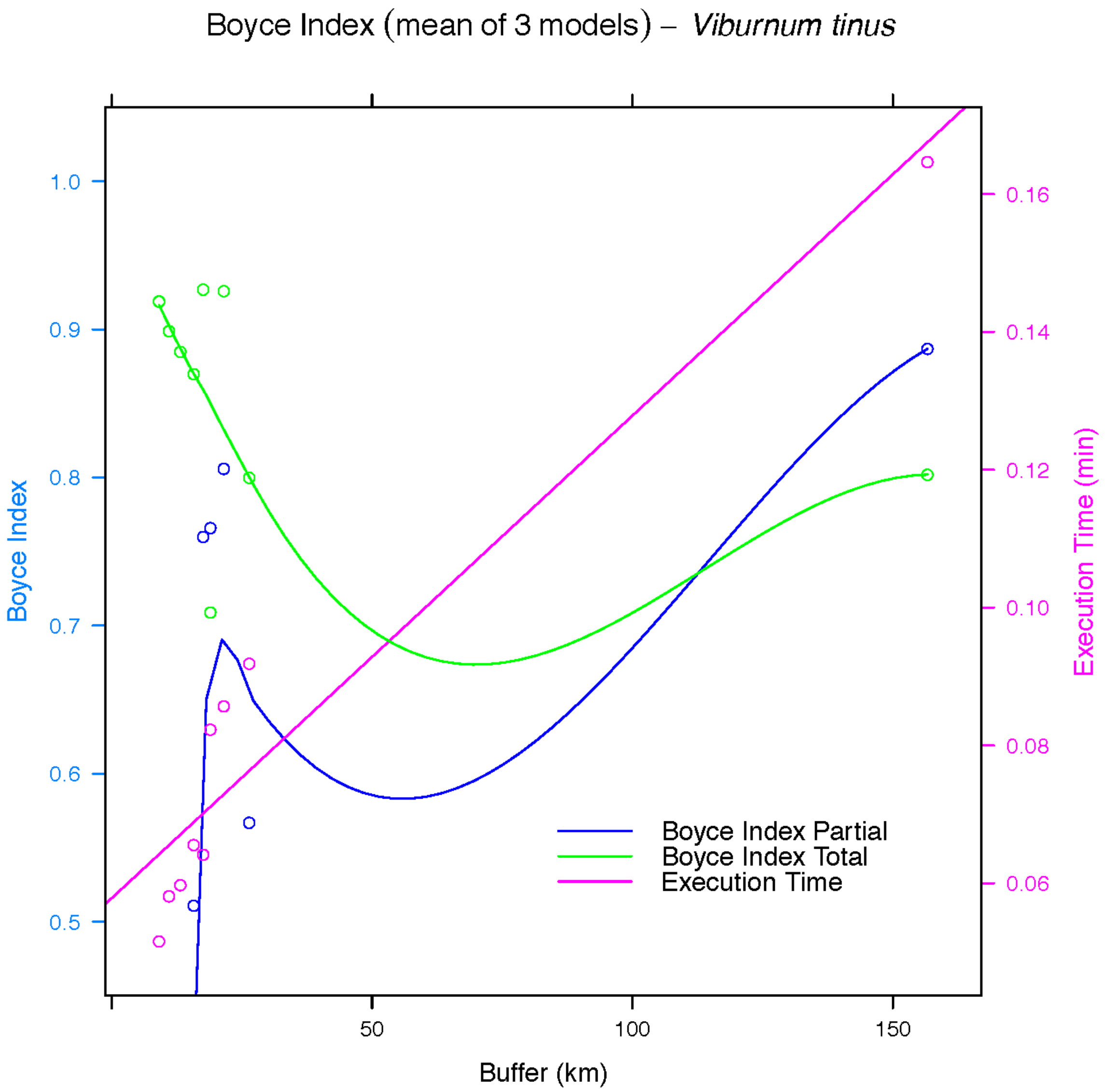

Boyce Index (mean of 3 models) – *Viburnum tinus*
Boyce Index
1.0
0.9
0.8
0.7
0.6
0.5
Execution Time (min)
0.16
0.14
0.12
0.10
0.08
0.06
Boyce Index Partial
Boyce Index Total
Execution Time
50
100
150
Buffer (km)


69
